# Cancer relevance of circulating antibodies against LINE-1 antigens in humans

**DOI:** 10.1101/2023.02.03.526997

**Authors:** Alexandra V. Vylegzhanina, Ivan A. Bespalov, Ksenia A. Novototskaya-Vlasova, Brandon M. Hall, Anatoli S. Gleiberman, Han Yu, Olga V. Leontieva, Katerina I. Leonova, Oleg V. Kurnasov, Andrei L. Osterman, Grace K. Dy, Alexey A. Komissarov, Elena Vasilieva, Jeff Gehlhausen, Akiko Iwasaki, Christine B. Ambrosone, Takemasa Tsuji, Junko Matsuzaki, Kunle Odunsi, Ekaterina L. Andrianova, Andrei V. Gudkov

## Abstract

LINE-1 (L1), the most abundant family of autonomous retrotransposons occupying over 17% of human DNA, is epigenetically silenced in normal tissues but frequently derepressed in cancer, suggesting that L1-encoded proteins may act as tumor-associated antigens recognized by the immune system. Here, we established an immunoassay for detecting circulating autoantibodies against L1 proteins in human blood. Using this assay in >3,000 individuals with or without cancer, we observed significantly higher IgG titers against L1-encoded ORF1p and ORF2p in patients with lung, pancreatic, ovarian, esophageal, and liver cancers compared to healthy individuals. Remarkably, elevated levels of anti-ORF1p-reactive IgG were observed in cancer patients with disease stages 1 and 2, indicating that immune response to L1 antigens can occur at early phases of carcinogenesis. We conclude that the antibody response against L1 antigens could contribute to the diagnosis and determination of immunoreactivity of tumors among cancer types that frequently escape early detection.

## Introduction

Long interspersed nuclear element-1 (LINE-1 or L1) is a non-long terminal repeat (LTR) retrotransposon that is comprised of hundreds of thousands of copies accumulated during mammalian evolution (Ostertag et al., 2001). In humans, about a half million copies of L1 collectively occupy over 17% of DNA (Lander et al., 2001). The vast majority of these sequences are functionally deficient due to truncations, internal rearrangements, and mutations. Only ∼150 copies of L1 elements are retrotransposition-competent in modern humans (Brouha et al., 2003). Among these full-length L1s, only a few “hot” loci contribute to the bulk of L1 mRNA expression (Kaul et al., 2020). Intact L1 sequences are approximately 6 kilobases in length and encode two polypeptides, open reading frame 1 (ORF1p) and 2 (ORF2p) (Dombroski et al., 1991). ORF1p is an RNA-binding protein (Holmes et al., 1992; Martin et al., 2001) and ORF2p is a multifunctional protein combining reverse transcriptase (RT) and endonuclease (integrase) activities essential for replication and expansion of L1 and other non-autonomous retrotransposons (Feng et al., 1996). The L1 replication machinery is believed to be responsible for most reverse transcription-driven integration events in the genome, including those involved in the expansion of short interspersed nuclear elements (SINE), pericentromeric satellite DNA, and processed pseudogenes (Ostertag et al., 2001; Kajikawa et al., 2002).

L1 elements are transcriptionally silenced in most normal cells due to multiple mechanisms of epigenetic repression, including DNA methylation and chromatin modifications; the latter mechanism involves histone deacetylases (e.g., Sirtuin 6 (Tasselli et al., 2017)), tumor suppressors p53 (Tiwari et al., 2020) and Rb (Montoya-Durango et al., 2010). Desilencing of L1 creates multiple deleterious risks for the cell and the organism (Zhang et al., 2020). In germline cells, it can lead to inherited diseases due to insertional mutagenesis (Wang et al., 2017; Bao et al., 2012). In somatic cells, it drives multiple mechanisms of genetic and epigenetic instability stemming from insertional mutagenesis (Rodriguez-Martin et al., 2020; Tubio et al., 2014), DNA damage caused by L1 ‘s endonuclease activity (Chen et al., 2005), and activation of interferon- and NF-κB-mediated inflammatory responses (Gazquez-Gutierrez et al., 2021; Chuong et al., 2016).

L1 elements are frequently derepressed in tumors, contributing to cancer genome instability (Zhang et al., 2020; Rodić et al., 2014) based on the inherent potential of somatic L1 insertions to drive tumorigenesis by activating oncogenes or inactivating tumor suppressor genes (Tiwari et al., 2020; Montoya-Durango et al., 2010). This situation is realized in a large proportion of human tumors that express L1 antigens (Zhang et al., 2020; Ardeljan et al., 2017) and experience L1 expansion in their genome (Chen et al., 2005; Beck et al., 2010). New L1 insertions have been identified in 53% of tumors sequenced (Rodriguez-Martin et al., 2020).

L1 activity differs between and within cancer types and may change during cancer progression (Rodić et al., 2013; Rodić et al., 2018). The most recent data defines esophageal adenocarcinoma and lung squamous cell carcinoma as having the highest rates of retrotransposition among human cancers. In a study analyzing 246 pancreatic cancer samples, approximately half exhibited an active retrotransposition process (Rodriguez-Martin et al., 2020). This proportion would likely be higher if SINE retrotranspositions were also considered. Based on the above, L1 ORF1p and ORF2p proteins have been considered as cancer biomarkers (Ardeljan et al., 2019; Sharp et al., 2020; Hosseinnejad et al., 2018). According to immunohistochemical staining for ORF1p, 47% of 1,027 samples representing more than 20 cancer types scored positive for L1 expression (Rodić et al., 2014). ORF2p was less frequently detected in tumors, which is not surprising given the many-fold lower expression of ORF2p compared to ORF1p (Dai et al., 2014). Within ORF2p-positive tumors, a switch from cytoplasmic to nuclear staining for ORF2p was reported during tumor progression (De Luca et al., 2016; Gualtieri et al., 2013).

Regardless of established L1 expression in multiple cancer types, it has not been developed a reliable assay for the detection of L1 antigens in blood circulation, and thus, they have not yet been translated into clinical biomarkers for cancer detection (Ardeljan et al., 2019). There are several possible explanations for this: being intracellular non-secreted proteins, L1 proteins may not enter circulation in the amounts sufficient for detection, may undergo rapid degradation, or may be masked by anti-ORF1p/ORF2p antibodies.

A similar situation is found regarding the development of genomic approaches to the detection of L1 activity. A substantial effort towards developing computational techniques for analyzing L1 sequences in genomic DNA has advanced our ability to detect frequent expansions and to map new copies of L1 in tumors (Gonzalez-Perez et al., 2013). The analysis of L1-derived sequences in liquid biopsies demonstrated a significant decrease in L1 methylation in cell-free DNA of cancer patients as compared to healthy controls. However, neither of these observations has been translated into clinically feasible diagnostic assays suitable for early cancer detection (Brenner et al., 2011; Gainetdinov et al., 2016).

During normal development, L1 expression is limited to early stages of embryogenesis (Garcia-Perez et al., 2007), brain neurons, especially in elderly subjects (Baillie et al., 2011; Upton et al., 2015; Evrony et al., 2015; Peze-Heidsieck et al., 2022), and specific cell subpopulations in the testis (Branciforte et al., 1994; Blythe et al., 2021; Ergün et al., 2004). Since these sites are all behind immunological barriers and, therefore, do not participate in the formation of immune tolerance to autoantigens, one would expect that L1 antigens expressed in tumors should be recognized by the immune system as tumor-associated antigens and induce an adaptive immune response. Based on this assumption, we hypothesized that the presence of L1-positive cancer could be associated with the development of autoantibody response to L1 antigens. A major possibility of such a response is supported by a report of anti-ORF1p antibodies in patients with autoimmune diseases (Carter et al., 2020). We expected that antibody detection may have clear advantages over detecting antigens themselves, since adaptive immunity can provide an amplified response to the presence of a biomarker that may have limited access to circulation and could be undetectable in the blood and because of long *in vivo* half-life of antibodies.

Driven by these considerations, we have developed a robust immunoassay, named ABLE (**<u>a</u>**nti**<u>b</u>**odies to **<u>L</u>**1 detected by **<u>E</u>**LISA), capable of specific semi-quantitative detection of circulating antibodies against L1 antigens in human sera. Using ABLE, we have analyzed >2,500 blood samples from patients with 14 cancer types along with >300 cancer-free individuals, either healthy or with cancer-unrelated health problems. The obtained results support our hypothesis, demonstrating a frequent association of the ongoing carcinogenic process with elevated levels of anti-ORF1p antibodies in blood. Remarkably, this association was observed in both early and advanced disease stages, supporting a potential utility of ABLE for cancer detection and determination of tumor immunoreactivity.

## Results

### Detection of anti-L1 antibodies in patients with a variety of cancer types and in healthy individuals

The hypothesis underlying the present study is schematically described in **Fig. 1A**. For detection of anti-ORF1p or anti-ORF2p antibodies in serum samples, we utilized an indirect ELISA technique using 96-well plates coated with recombinant polypeptides representing L1 antigens: a full-length ORF1p or a 47-kDa fragment of ORF2p covering the region spanning 367-771 amino acids (see Materials and Methods). We used the full-length human L1 ORF1p sequence with no modifications, since, according to a BLAST search, ORF1p has no significant similarities with other human proteins that could result in cross-reactivity of anti-ORF1p antibodies. The ELISA protocol was optimized to increase sensitivity and reduce background.

**Figure 1.**
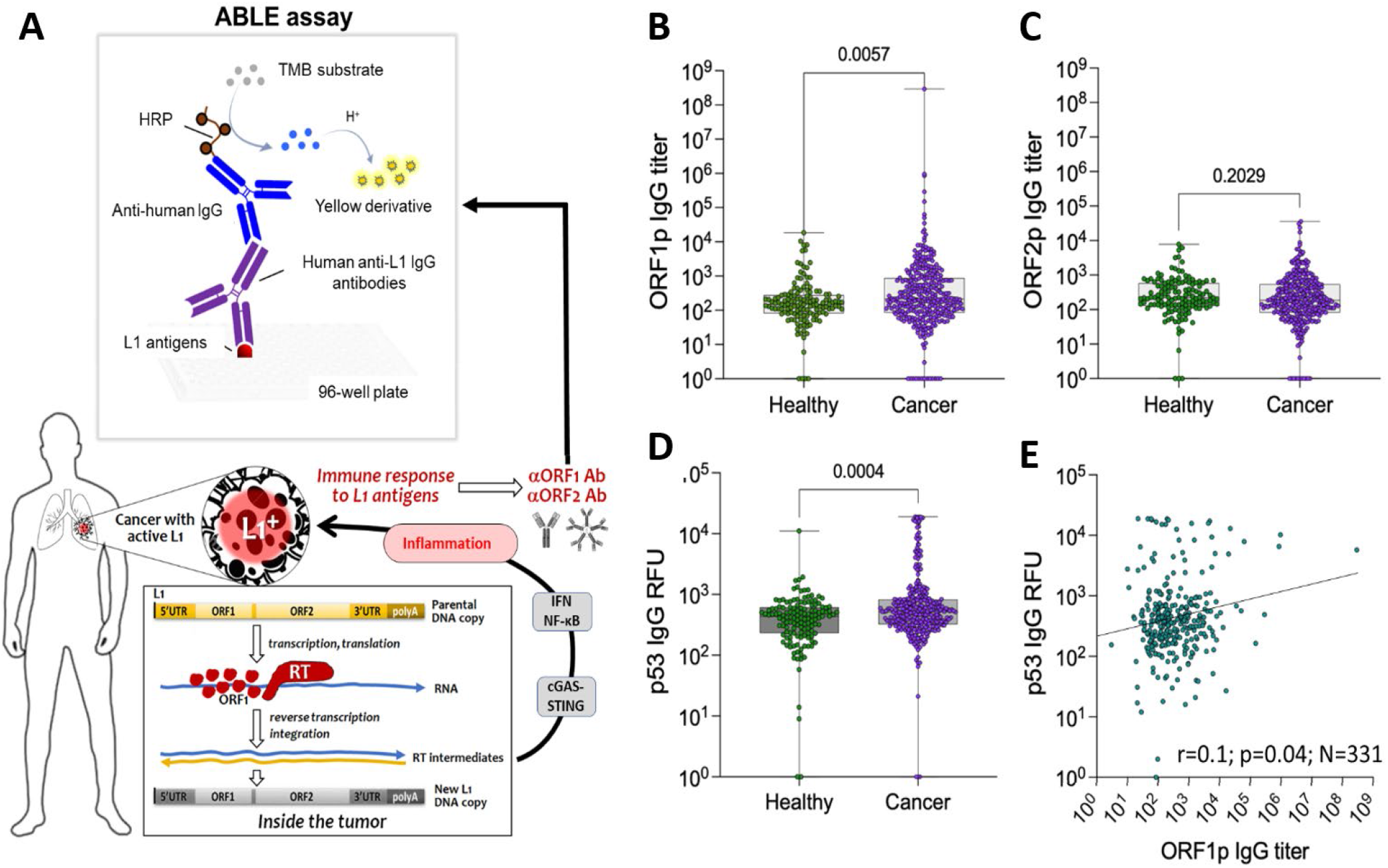
Approach and proof of concept. **A**. Hypotheses: proteins encoded by L1 are expected to be recognized as tumor-associated antigens and become targets for antibody response. This response is stimulated by the cGAS-STING-mediated induction of inflammation. Circulating Abs to L1 antigens can be used as cancer biomarkers detected by ABLE (antibodies to L1 detected by ELISA) assay. **B, C, E**. Comparison of antibody levels in healthy individuals (N=137) and cancer patients (N=331) in the assays for anti-ORF1p (**B**), anti-ORF2p (**C**) and p53 (**D**). The significance of differences is assessed by Mann-Whitney U-test. **E**. Spearman ‘s Correlation between anti-ORF1p IgG titers and anti-p53 IgG signals in serum samples of cancer patients (N=331). All p-value less than 0.05 is considered significant.

To test the existence of the principal phenomenon (i.e., development of an antibody response to L1 antigens in cancer patients), at first we analyzed a mixed set of 331 serum samples representing 14 solid tumors, including: ovarian (N=24), breast (N=24), lung (N=24), colorectal (N=24), esophageal (N=24), renal (N=24), liver (N=24), pancreatic (N=24), prostatic (N=24), gastric (N=20), and bladder (N=24) cancers, soft tissue sarcoma (N=24), melanoma (N=23), and glioblastoma (N=24), along with 137 healthy individuals. The titers of anti-ORF1p IgG antibodies were found to be significantly higher in the cancer patient population compared to healthy subjects (**Fig. 1B**). The small sample sizes for each cancer type in this study limited our ability to reliably classify which cancer types exhibit elevated anti-L1 antibodies (**Fig. S1A**).

Evaluating anti-ORF2p IgG titers from the same samples did not show a significant difference between patients with cancer compared to healthy individuals (**Fig. 1C, Fig. S1B**). Furthermore, maximal anti-ORF2p IgG titers were more than two orders of magnitude lower that the maximal titers of anti-ORF1p antibodies.

The same serum samples were also used to detect IgG against p53, a tumor suppressor protein most frequently mutated in cancer, to which autoantibodies have been considered as a cancer detection biomarker (Liu et al., 2020). The rationale for comparison of humoral immune responses to L1 antigens and p53 is based on the involvement of wild-type p53 in epigenetic repression of L1 (Wylie et al.,2016) and the fact that p53 mutation correlated with L1 ORF1p expression in some cancer types (McKerrow et al., 2022). As with antibody response to ORF1p, the same cancer patients showed significantly higher elevation in anti-p53 IgG (**Fig. 1D, Fig. S1C**). A weak but significant positive correlation between anti-ORF1p and anti-p53 IgG signals was observed (Spearman ‘s correlation: r=0.1, p=0.04; **Fig. 1E**).

To assess specificity of the detected antibodies to L1 proteins, we analyzed serum samples from four ovarian cancer patients that had 10^7^-10^8^ anti-ORF1p IgG titers in the ABLE as the source of antibodies for immunofluorescent staining and immunoblot analysis of HeLa cells expressing tetracycline-inducible human L1 (see Materials and Methods, **Fig. S2**). Analysis revealed bright immunofluorescent staining of HeLa cells with induced L1 expression (but not the same cells without L1 induction) (**Fig. 2A**) and a single band with the expected size for ORF1p (40-kDa) on the immunoblots of the lysates of HeLa cells after doxycycline-induced L1 expression (**Fig. 2B**). These results were identical to those obtained with rabbit antibodies to ORF1p used as control (**Fig. 2A&B**, left panels).

**Figure 2.**
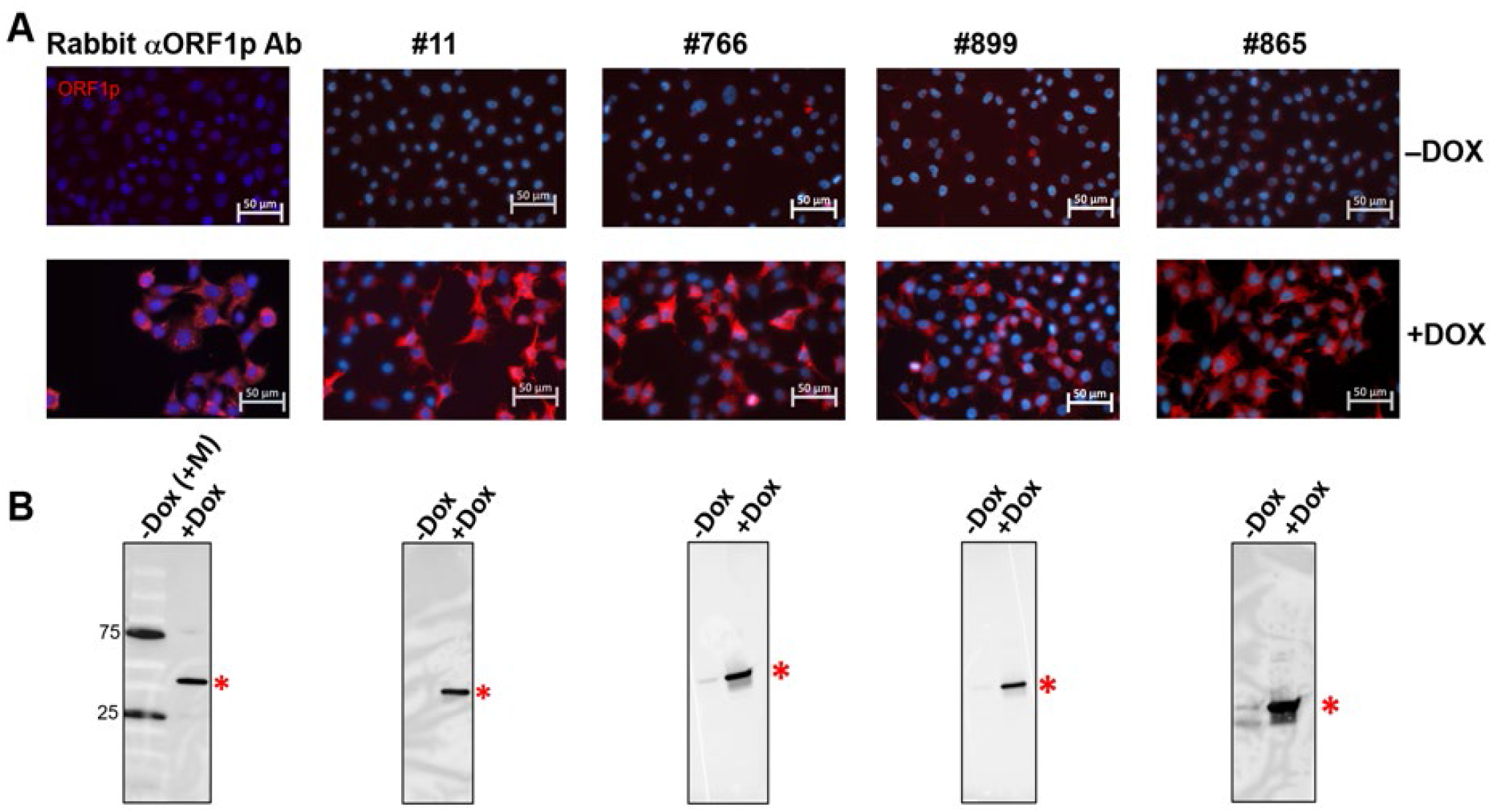
Relevance of antibodies detected by ABLE to L1 ORF1p. **A**. The results of immunofluorescent staining and **B**. Immunoblot analysis of HeLa cells transduced with Tet-inducible L1/Gluc reporter construct with (+DOX) and without (-DOX) L1 induction by doxycycline (**Fig. S2**). “Rabbit αORF1p Ab” – polyclonal anti-ORF1p rabbit antiserum. The other panels represent staining with 50,000-fold diluted serum samples from ovarian cancer patients with 107-108 anti-ORF1p IgG titers in ABLEORF1 assay. “M” stands for the protein size marker added to the sample along with HeLa cells lysate without DOX L1 induction. The asterisks on the lower panel correspond to ORF1p standard (40-kDa) band.

Sera that were negative in the ABLE (titers below 450) failed to produce signals in both immunofluorescent staining and immunoblots (**Table S1**).

Within the serum samples analyzed, a clear relevance of anti-ORF1p IgG titers in ABLE with sample positivity in immunofluorescent staining of cells with induced L1 and immunoblot-based detection of ORF1p was observed (**Table S1, Fig. S3**). Rare inconsistencies, where ABLE-positive serum sample failed to detect ORF1p signal in HeLa cells by immunostaining, could be explained by the specificity of antibodies in these samples to conformational ORF1p epitopes.

The fact that patients ‘ serum samples used in different dilutions for immunofluorescent and immunoblot staining demonstrated highly specific recognition of ORF1p among numerous protein present in fixed human cells or human cell lysates indicates that anti-ORF1p IgG appeared as a major autoimmune-reactive antibodies in circulation in most subjects that had 10^4^-10^9^ anti-ORF1p IgG titers in ABLE (**Fig. 2B, Fig. S3**).

We were unable to detect ORF2p using immunofluorescent or immunoblot staining, even with those serum samples that had 10^4^-10^5^ anti-ORF2p titers in ABLE. This is consistent with overall lower Ab levels to this L1 protein but may also be associated with conformational ORF2p epitopes.

To test whether antibody response to L1 antigens is associated with the presence of L1 antigens in circulation, we established a sandwich ELISA (see Materials and Methods) capable of detecting 0.1 ng/mL of ORF1p and used it to analyze the same set of serum samples from cancer patients. The presence of high antibody levels against ORF1p in circulation could mask the signal in this assay (**Fig. S4A**). To address this possibility, we introduced a protein-antibody denaturing step in the protocol that indeed increased the assay sensitivity in the presence of anti-ORF1p antibodies in model experiments (**Fig. S4B**). Only 3% of tested samples possessed detectable concentrations of ORF1p ranging between 0.12 and 5.5 ng/ml (median 0.36 ng/mL; N=331 cancer patient samples and N=72 healthy individuals) and the denaturation step in the sandwich ELISA did not generate any additional signals. Consistently, these 3% samples did not belong to those with the highest anti-ORF1p IgG titers. These observations indicate the advantage of anti-L1 ORF1p IgG vs ORF1p antigen measurement in the blood, at least within the current limit of detection of the ORF1p sandwich ELISA.

### Disease stage relevance of anti-ORF1p IgG titers

We expanded our study to several cancer types with poor prognoses and the treatment of which would greatly benefit from early detection. Specifically, we analyzed an additional set of 2,484 serum samples from patients with ovarian (N=979), esophageal (N=377), lung (N=907), pancreatic (N=124) and liver cancers (N=217) that included patients with early (1-2) and advanced (3-4) disease stages (**Table S6**). The rationale for this selection is illustrated in **Fig. 3A**. The five cancer types selected are characterized by a poor five-year survival, and four of them (esophageal, pancreatic, liver and ovarian) currently do not have clinically-approved screening protocols in people who are at average risk. The screening procedure currently used for the lung cancer detection in a high-risk population is known to be insufficiently accurate and involves an invasive biopsy collection step (Seijo et al., 2019). These cancers (with exception of hepatocellular carcinoma) have a high frequency of L1 derepression, as indicated by immunohistochemical assessment and the detection of retrotranspositions found in earlier reports (Ardeljan et al., 2017; Rodriguez-Martin et al., 2020).

**Figure 3.**
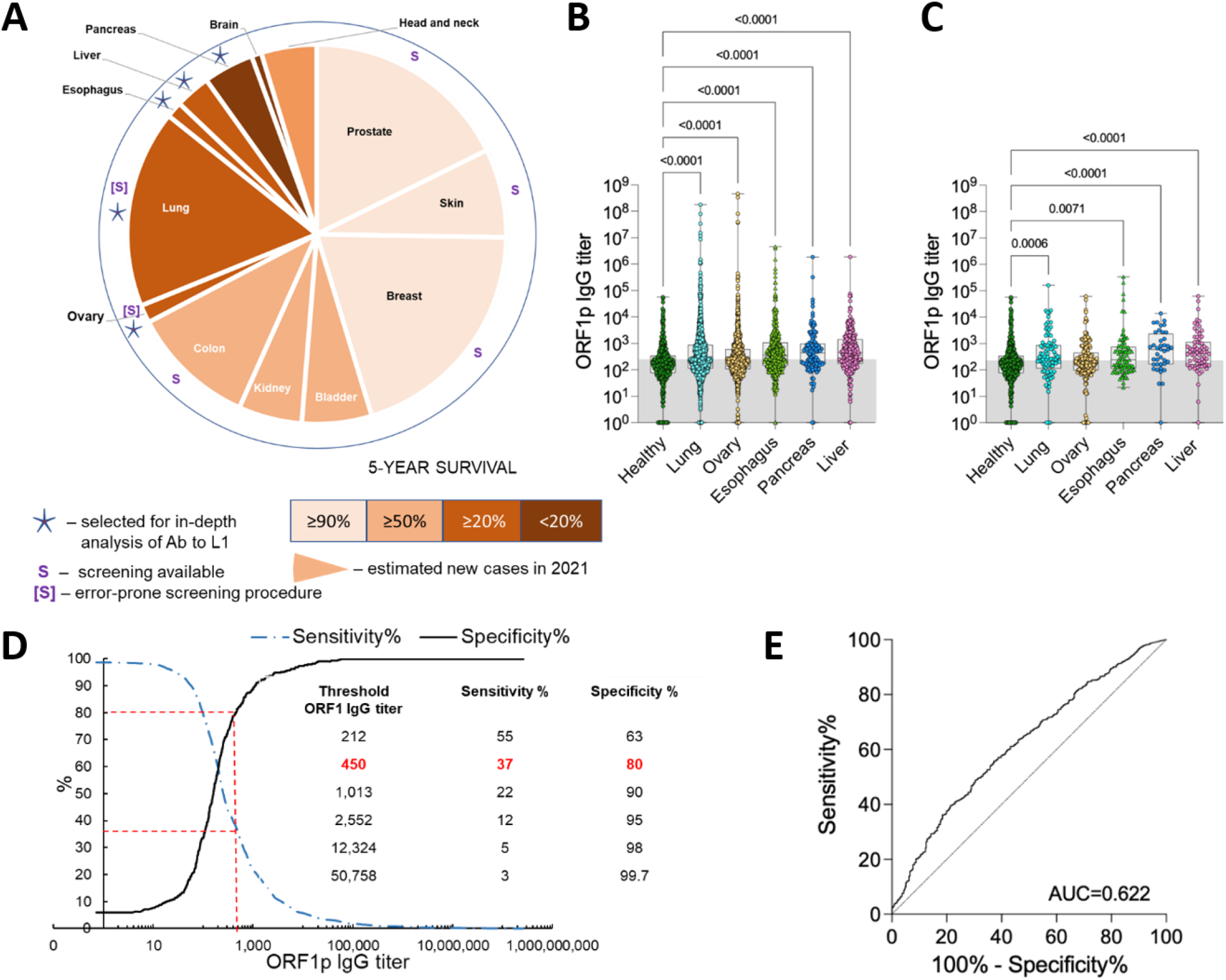
Anti-ORF1p IgG response in serum samples from patients with five cancer types. **A**. Selection of cancers for ABLE analysis. Sector sizes are proportional to the incidence of these cancers in the US in 2021 (https://seer.cancer.gov/statfacts/). Color code indicates a 5-year survival. “S” stands for the existence of approved cancer screening protocols. **B**. Detection of anti-ORF1p IgG titers in serum samples of patients with the indicated cancer types of all stages. Sample size: ovary (N=979), esophagus (N=377), lung (N=907), pancreas (N=124), liver (N=217) cancers and healthy (N=352). **C**. The same as panel B; only samples from patients with cancer stages 1 and 2 are shown. Sample size: ovary (N=193), esophagus (N=79), lung (N=90), pancreas (N=40) and liver (N=67) cancers and healthy (N=352). The grey areas in panels B and C mark samples below the anti-ORF1p IgG titer threshold of 450. Statistics were calculated by Dunn ‘s multiple comparison test with adjusted p-value for anti-ORF1p IgG titers. All p-value less than 0.05 is considered significant. **D**. ABLEORF1 assay threshold determination. **E**. The ROC curves indicating specificity and sensitivity of ABLEORF1 assay for combined sample set of 5 cancers (ovary (N=979), esophagus (N=377), lung (N=907), pancreas (N=124), liver (N=217) cancers and healthy (N=352)).

The comparison with 352 samples from healthy individuals revealed significantly elevated anti-ORF1p IgG titers in all five selected cancers as judged by non-parametric Dunn ‘s test with adjusted p-value (**Fig. 3B**). Importantly, these differences retain high significance if we limit the analysis to patients with early (1-2) stages of lung (N=90), esophageal (N=79), pancreatic (N=40) and liver (N=67) cancers even though the number of cases in each category became substantially smaller (**Fig. 3C**).

The analysis of anti-ORF2p IgG showed similar, though less pronounced, effects. Significantly elevated IgG titers to the ORF2p were found for three out of five selected cancer categories (ovarian, pancreatic, and liver cancer) (**Fig. S5A**). We found positive correlation between anti-ORF1p and ORF2p IgG titers in all these cancers (**Fig. S6**). As far as the early-stage cancers is concerned, increased anti-ORF2p IgG titers compared to healthy population was found for stages 1-2 of pancreatic and liver cancers (**Fig. S5B**) opening the opportunity for a more precise early detection of these cancer types if ABLEs for ORF1p and ORF2p are combined.

The difference between cancer patients and cancer-free subjects is evident in the subpopulation with elevated levels of anti-ORF1p IgG and the higher this elevation is, the stronger the difference (**Fig. 3D**), thus increasing cancer predictive power of the signal. Receiver Operating Characteristic (ROC) curves were built to estimate anti-ORF1p IgG titer thresholds in ABLE providing an acceptable combination of specificity (80%) and sensitivity (37%) (**Fig. 3D-E, Fig. S12**). From the combined ROC curve an average titer threshold for 5 cancer types was 450. In this case the use of ABLE assay for cancer diagnostics is complicated by a frequent presence of detectable levels of circulating IgG to L1 antigens in healthy humans (∼20% of healthy individuals had anti-ORF1p IgG titers above threshold 450). Therefore, this parameter cannot be used as an indicator of the lack of carcinogenic process but individuals with a positive test, with anti-ORF1p IgG titers, for example, more than 50,758, have only 0.3% chance to be free from cancer.

### Testing factors and conditions potentially confounding cancer relevance of ABLE results

Recent reports on L1 derepression in senescent cells (De Cecco et al., 2019) suggested a potential age dependence of antibody response to L1 antigens. Indeed, we found an increase of anti-ORF1p IgG titers with age (Spearman’s correlation in healthy: r=0.2, p=0.0001; N=352; 20-89 years of age (**Fig. S7A**) and cancer: r=0.08; p=0.0001; N=2,470; 13-95 years of age (**Fig. S7B**)), rather weak but statistically significant. Meanwhile, logistic regression analysis established the significant (p=0.03) association between cancer and anti-ORF1p IgG titers after adjustment for age (cancer (N=2,470) and healthy (N=352); 13-95 years of age) (**Table S2**). So, we concluded that anti-ORF1p IgG titer’s association with cancer is not confounded by age.

No racial differences in anti-ORF1p IgG titers were found in healthy or cancer population, as analyzed via Dunn’s multiple comparison test (**Table S3**). Gender-specific differences were observed only in lung cancer samples, where anti-ORF1p IgG titers were observed to be higher in male patients compared to female (p=0.004; Mann-Whitney U-test) (**Table S4**). No significant differences were found in other cancer types analyzed. No dependence of anti-ORF1p IgG titers on smoking history was found in lung cancer patients (**Fig. S8A**) or in a combined set of pancreatic, esophageal, and liver cancer (**Fig. S8B**).

Similarly, we did not find any significant differences of anti-ORF1p IgG titers in a cross-sectional study of cancer patients treated by radiation therapy, immunotherapy, or chemotherapy compared to non-treated cancer patients (**Fig. S9**). However, our analysis included only patients whose disease advanced following treatments and did not include full responders. Whether anti-ORF1p IgG titers decrease following cancer eradications remains to be determined.

We also tested the stability of anti-ORF1p IgG levels in a longitudinal study of 18 healthy individuals whose blood samples were collected three times (days 0, 49 and 189) within 6 months. The levels of anti-ORF1p IgG titers ranged substantially between individuals, but these levels remained stable within the observation period (**Fig. S10**).

L1 expression can trigger inflammatory pathways mediated by cGAS-STING signaling (De Cecco et al., 2019; Simon et al., 2019). Therefore, it would be important to assess potential relevance of antibodies against L1 antigens within inflammation-associated diseases. The titers of IgG against L1 antigens were evaluated via the ABLE in serum samples from patients with recent myocardial infarction (MI, N=37), as well as from patients with chronic obstructive pulmonary disease and/or pneumonia (COPD/PN, N=33). We found no statistically significant differences compared to the healthy population, suggesting that COPD/PN and MI does not influence the levels of IgG against L1 antigens (**Table S5**).

Thus, we conclude that anti-ORF1p IgG titers are a cancer-associated parameter that maintains high stability over time, is not confounded by age and gender, race, smoking, inflammation, nor cancer treatment history.

Prior publication reported the presence of anti-ORF1p antibodies in patients with systemic lupus erythematosus (SLE) (Antiochos et al., 2022; Carter et al., 2020). Here we analyzed a set of 17 samples from patients with SLE and found elevated anti-ORF1p (but not anti-ORF2p) IgG titers as compared to healthy individuals (**Fig. S11A-B**). These results confirm that development of anti-ORF1p IgG response might be common among patients with autoimmune diseases and would need to be considered during analysis. It is noteworthy that anti-p53 IgG signals in SLE patients were significantly higher than in healthy subjects (**Fig. S11C**).

## Discussion

We developed an immunoassay named ABLE for the detection of circulating antibodies against L1-encoded proteins and used it for the analysis of >3,000 blood samples that included healthy individuals and patients with a variety of cancer, autoimmune, cardiovascular and lung diseases. We found that IgG antibodies against ORF1p, and to a lesser extend ORF2p, are frequently detected in humans, indicating that that L1 antigens are commonly recognized by the immune system and induce humoral immune responses. Among the disease categories tested, cancer showed the strongest elevation of serum IgG antibody levels to L1 antigens. Extended testing validated this finding in patients with lung (NSCL, SCL), esophageal, ovarian, pancreatic, and liver cancers; other cancer types remain to be thoroughly analyzed. L1 desilencing was previously considered as a cancer biomarker detectable by hypomethylation in circulating cell-free DNA in liquid biopsies (Ardeljan et al., 2017; Gezer et al., 2022). Here, we demonstrate that the epigenetic dysregulation of L1 can generate an additional type of biomarker – circulating autoantibodies against L1 antigens – that can be detected by a simple and robust blood test. Antibodies against L1 antigens in the blood are detected more readily than the antigens themselves, owing to the amplification of the L1 expression signal by the immune response. The higher IgG titers observed for ORF1p over ORF2p likely reflects the many-fold higher expression of ORF1p compared to ORF2p (Taylor et al., 2013; Dai et al., 2014; Ardeljan et al., 2019).

Another valuable property of anti-L1 IgG antibodies with respect to their diagnostic potential is that their elevation is frequently observed in patients with early (1-2) stages of cancer. This is likely explained by the fact that L1 expression frequently occurs at early stages of malignant transformation and either stays high during tumor progression (Zhang et al., 2020 and Grundy et al., 2022; Kelsey et al., 2021) or, in some instances, may decrease (Guler et al., 2017). This feature distinguishes anti-L1 antibody response from that of other autoantibodies considered as potential cancer biomarkers, which are limited in their value for early cancer detection since they are more commonly associated with advanced stages of disease (e.g., to p53, heat shock proteins, etc. (Shi et al., 2017 ; Liu et al., 2020; de Jonge et al., 2021)).

Only a portion of cancer patients have elevated levels of anti-L1 IgG antibodies, which precludes the use of the ABLE as a negative screening test for cancer. However, the potential utility of the ABLE is found in strong immune response in patients with cancer that have an abnormally high elevation of anti-L1 IgG titers.

This apparent advantage of anti-L1 antibodies, however, is confounded by their infrequent appearance in healthy volunteers without known cancers or autoimmune disease. A possible explanation for the presence of detectable levels of anti-ORF1p antibodies in healthy subjects is the presence of occult disease, or alternatively, that these antibodies remain as a memory of adaptive immune response to past events associated with the derepression of L1 (e.g., initiated carcinogenic processes intercepted by anticancer defense mechanisms). In fact, it has been demonstrated that L1 derepression can precede cancer development (Kelsey et al., 2021). According to this model, low levels of anti-L1 antibodies in healthy individuals may be “remnants” of a past success of antitumor immunity, whereas the strong accumulation of anti-L1 antibody titers in the blood likely signals persistent activation of associated memory B-cells by the presence of L1-expressing cells and may reflect an inability of the immune system to effectively eradicate these cells (**Fig. 4**).

**Figure 4.**
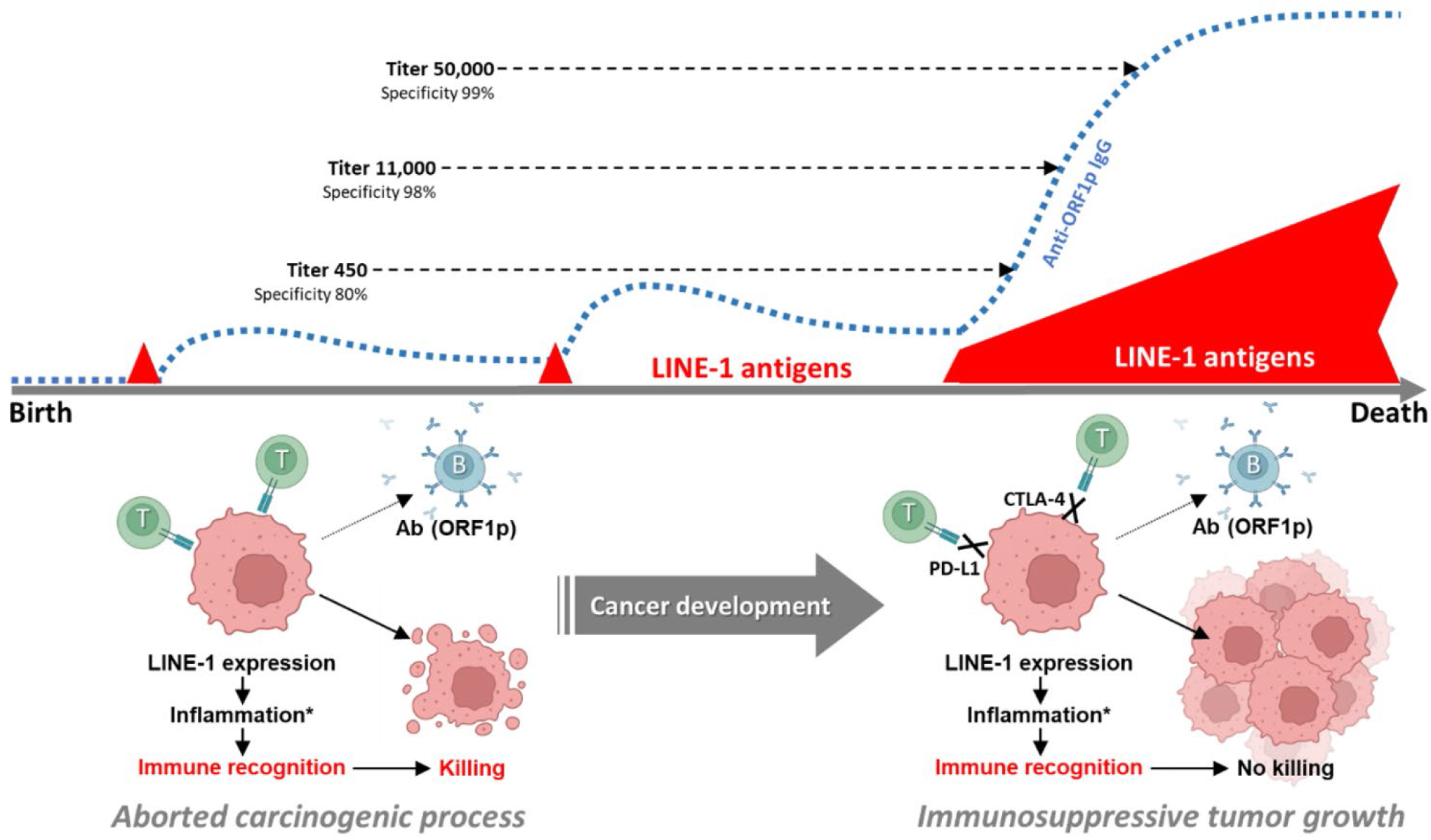
Schematic description of a hypothetical relationship between circulating IgG titers to LINE-1 antigens and cancer susceptibility to immunotherapy by immune checkpoint inhibitors. Presence of low titers of anti-LINE-1 IgG in healthy subjects can be explained by immune response that occurred to precancerous lesions associated with derepression of LINE-1 that were aborted due to effective T-cell response. These events generate memory B-cells and the presence of low titers of anti-LINE-1 antibodies even in a cancer-free organism. Development of cancer that passivate T-cell response due to the engagement of immune checkpoint factors PD-L1 and CTLA-4 becomes a growing source of LINE-1 antigens. If this cancer expresses LINE-1 and retains the ability to be recognized by the immune system (e.g., the ability to present antigens), it will then induce elevation of anti-LINE1 IgG levels, which will signal about principal immunoreactivity of the tumor. *This response is stimulated by the cGAS-STING-mediated induction of inflammation

Another apparent limitation of ABLE is its inability to distinguish among multiple cancer types. In fact, similar frequencies of elevated IgG levels against ORF1p were observed in patients with five cancer types and this number will likely increase after other cancer types are analyzed. These constrains do not allow us to view ABLE as a stand-alone screening test for cancer presence but rather a supplementing assay to be included as a component in already used or emerging cancer diagnostic panels in populations of high risk, such as, for example, CA125 in ovarian cancer patients (Scholler et al., 2007) or different versions of liquid biopsy-based assays (Yee et al., 2020).

p53 was shown to play a role in epigenetic repression of L1 (Wylie et al., 2016). In fact, a recent study demonstrated that L1 expression in cancer correlates with p53 mutations (McKerrow et al., 2022). Therefore, it would be reasonable to expect a correlation between p53 inactivation in the tumor and the development of anti-L1 antibody response. We could not directly address this possibility in our study since we do not have the information on the p53-status of tumors from patients whose serum samples we analyzed. However, we could test autoantibodies to p53 in our panel knowing that the development of anti-p53 antibodies usually follows p53 mutations that create highly stable, aberrantly accumulated, and therefore immunogenic, protein. It was shown that level of anti-p53 IgG was higher among cancer serum samples with elevated anti-ORF1p IgG (**Fig. 1E**). The immunogenicity of L1 autoantigens may point at the possibility that their effective recognition could be part of evolutionary developed protective mechanisms against L1 activation.

These considerations suggest another potential clinical application for ABLE: its potential to distinguish between the tumors susceptible and resistant to immunotherapy with the inhibitors of immunological checkpoint factors PD-1 or CTLA-4 that are currently approved for the treatment of multiple cancers (**Fig. 4**) (Seidel et al., 2018; Vaddepally et al., 2020). The lack of reliable predictive tests for defining subsets of patients capable of responding to immunotherapy is a significant clinical problem (Liberini et al., 2021; Filipovic et al., 2020). In the context of our hypothesis, high levels of anti-L1 IgG mean that the immune system “senses” the tumor but cannot effectively eradicate it, suggesting the potential for a high probability of success utilizing immunotherapy by immune checkpoint inhibitors. The potential impact of derepressed retrotransposons on sensitizing tumors to immune checkpoint inhibitor immunotherapy has been proposed (Strick et al., 2015; Rooney et al., 2015) and may reflect, in part, the sensitizing effects of DNA hypomethylating agents that are known to stimulate retrotransposon expression (Roulois et al., 2015 ; Leonova et al., 2013). Alternatively, low levels of anti-L1 antibodies in patients with advanced L1-positive cancer may be evidence of low tumor immunogenicity and may be predictive of a poor performance of immunotherapy. Whereas T-lymphocyte responses play a major role in tumor eradication, antibodies to neoantigens are usually harmless to the tumor and, therefore, may serve as long-lasting biomarkers of immune recognition that was unable to be translated into an antitumor effect. This hypothesis implies an association between B- and T-cell responses to the tumor. The utility of ABLE for this hypothetical indication could be tested by evaluating the association of anti-L1 antibodies with the response to immunotherapy in NSCL and SCL cancers - a subject of our future study.

There are several intriguing mechanistic questions to be addressed regarding the described observations. What is the source of L1 antigens that facilitate their exposure to the immune system? Are antibody responses to L1 a reflection of recognition of new antigens or they have some currently unknown function in control of L1 activity? The establishment of the ABLE assay described in this work makes it possible to address these and other questions arising from this exciting phenomenon.

## Acknowledgements

We thank Annmarie Nowak, Maria Tokuyama (Department of Microbiology and Immunology, Life Sciences Institute, The University of British Columbia, Vancouver, BC, Canada) and Insoo Kang (Department of Internal Medicine, Yale University) for their help with sample collection and processing. Most of data and samples for this study were provided by the Data Bank and BioRepository (DBBR), which is funded by the National Cancer Institute (NCI P30CA16056) and is a Roswell Park Comprehensive Cancer Center Support Grant shared resource. This work was supported by a contract from Genome Protection, Inc. to A.V.G.

## Author contributions

AVG, ELA and AVV designed and coordinated the study. KAN-V, IAB, ASG, BMH, OVL, KIL, AAK conducted the study. AVG, ELA, AVV, KVN-V, KIL, HU, KO, EV, AI analyzed and interpreted the data. KA, KO, GKD, EV, AI, JM, TT provided patient samples and data. ALO and OVK generated recombinant L1 antigens. AVG, AVV and ELA wrote the manuscript. All co-authors edited and approved the manuscript.

## Materials and Methods

### Study patients and sample collection

Serum samples from cancer patients were obtained from the Roswell Park Data Bank and BioRepository (DBBR), a CCSG Shared Resource which prospectively consents new patients to provide blood samples, to complete an epidemiological risk factor questionnaire, and for linkage of samples to clinical data and to tumor tissue samples. Newly diagnosed patients are consented prior to cancer treatment to investigators with IRB-approved protocols for collection of blood samples, completion of a risk factor questionnaire, and permission to link their samples to tumor tissue and to clinical data. DBBR specimens are rigorously collected, processed, and stored so that the integrity of potential analytes will be consistent across all patients and controls. Blood is drawn into tubes that are immediately sent to the DBBR laboratory for standardized preparation of serum, plasma, red blood cells, buffy coat, and DNA; per SOP, aliquots are frozen within one hour of collection and stored in liquid nitrogen to minimize sample degradation. Samples were collected between 2003 and 2021, cancer patients were included if they were a newly diagnosed patient with a blood sample drawn and banked in the DBBR or Ovarian Bank with information in the Cancer Registry.

In addition to enrolling cancer patients, the DBBR also uses a number of approaches to enroll men and women with no personal history of cancer. These include controls recruited at community events, visitors accompanying cancer patients and employee volunteers. Samples are collected, processed and stored using the same protocols as those used for cancer patients.

The sample set used in this study consists of 2,815 well-annotated samples from the DBBR and Ovarian Bank of RPCCC (female N=1,663, male N=1152) with the age range varying between 13 and 95 years old. It includes 14 cancer categories classified according the organ of cancer origin: 24 samples from glioblastoma brain tumor; 24 samples from breast cancer (infiltrating ductal carcinoma and infiltrating lobular carcinoma); 907 samples of lung cancer (non-small cell carcinoma and small cell carcinoma), 24 samples from colorectal adenocarcinoma, 24 samples of soft tissue sarcoma, 377 samples from esophageal adenocarcinoma and squamous cell carcinoma, 24 samples of renal cell carcinoma; 217 samples of hepatocellular carcinoma; 979 samples of serous ovarian carcinoma, 124 samples of pancreatic adenocarcinoma; 24 samples of prostatic adenocarcinoma; 23 samples of skin malignant melanoma, 20 samples from gastric adenocarcinoma, 24 samples of urinary bladder cancer (papillary urothelial carcinoma and transitional cell carcinoma).

We also included 78 samples from DBBR controls and 200 from healthy volunteers obtained from Innovation Research (https://www.innov-research.com/), as well as samples from I.V. Davydovsky Clinical City Hospital (Moscow Department of Healthcare, Moscow, Russia collected September 2021-April 2022. The study was approved by the Moscow City Ethics Committee of the Research Institute of the Organization of Health and Healthcare Management (protocol #LABS022021) and performed according to the Helsinki Declaration. These included samples from healthy volunteers (N=74), non-cancer patients with recent myocardial infarction (N=37) and patients with pulmonary diseases, chronic obstructive pulmonary disease and/or pneumonia (N=33). A total non-cancer dataset consists of 352 serum samples from healthy subjects; among them: female N=194, male N=158 with the age ranging between 20 and 89 years old.

### Blood samples from patients with autoimmune diseases

SLE patient samples were utilized from a repository generated in a previous study (Tokuyama et al., 2021) under a protocol approved by the Institutional Review Committee of Yale University (HIC #0303025105). All enrolled SLE patients were diagnosed according to American College of Rheumatology 1997 criteria. Plasma samples were obtained after processing peripheral blood from sodium-heparin coated tubes and stored as aliquots at –80°C.

### Recombinant L1 antigens

The protein-coding DNA sequence corresponding to human L1 ORF1p described in Uniport Entry Q9UN81 (LORF1_HUMAN) was custom-synthesized with codon optimization for expression in E. coli (GenScript Biotech) and cloned into the modified pET-49b(+) expression vector (Novagen) with a C-terminal His6-tag under control of the T7 promoter. Overexpression in E. coli strain T7 Express Iq (NEB, Cat. # C2833H) was optimized to yield ∼100 mg/L of recombinant ORF1p-His6 (346 aa; 41.1 kDa) in the form of inclusion bodies (IB). Cells harvested by centrifugation were resuspended in lysis buffer (20 mM HEPES pH 7.0; 0.5M NaCl; 0.5% Brij-35) and lysed by sonication. Insoluble fraction collected after centrifugation of the crude lysate was extracted with 7M Urea-based extraction/ capturing buffer (50mM Tris-HCl, pH 8.5; 250mM NaCl; 7M Urea; 20mM Imidazole; 0.15% Brij35; 2.5mM β-mercaptoethanol; 1/20 volume of expression culture). Capturing of ORF1p-His6 protein was performed using Immobilized-Metal Affinity Chromatography (IMAC) by loading of 7M Urea Extract on 5 mL HisTrap FF FPLC column (GE Healthcare) at 0.5 mL/minute flow rate followed by washing with 10 column volumes (CV) of the same 7M Urea-based buffer. Partial in-column refolding by washing with 10 CV of 4M Urea-based buffer with 0.5M NaCl and 0.3% Brij35 was followed by gradient elution with 20-300mM imidazole in 4M Urea-based buffer. Pooling of the main peak fractions yielded highly purified ORF1p protein (∼90% by SDS-PAGE analysis) without appreciable host cell proteins (HCPs) impurities. Two-step dialysis assisted transferring of recombinant ORF1p protein to storage buffer. Initial dialysis against storage buffer containing 2M Urea was followed by dialysis against storage buffer: 50mM Tris-HCl, pH 8.0; 0.5M NaCl; 2mM DTT; 0.05% Tween80; 5% Glycerol ‘. Sterile-filtered final protein sample with concentration adjusted to 1mg/ml was flash-frozen in aliquots in low protein binding (LPB) plastic vials before storage at -80°C. The protein-coding DNA sequence corresponding to fragment 367-771 of human L1 ORF2p described in Uniport Entry O00370 (LORF2_HUMAN); or as human NAG13 protein (GenBank ID: AAG27485.1) was custom-synthesized with codon optimization for expression in E. coli (GenScript Biotech). Cloning and expression of 367-771 fragment (413 aa; 47.6 kDa) of human L1 ORF2p was performed according to the same protocol as described above for human ORF1p, with exception of using the Size-Exclusion Chromatography (SEC) as final purification step after IMAC purification/refolding. HiLoad 16/600 Superdex-200 FPLC column (GE Healthcare) was equilibrated with 50mM Tris-HCl, pH 8.5; 2M Urea; 0.5M NaCl; 0.15% Brij35; 2.5mM DTT. SEC-eluted samples were subject to buffer exchange on Fast-Desalt FPLC column HiPrep 26/10 (GE Healthcare) in 50mM Tris-HCl, pH 8.0; 0.3M NaCl; 0.5mM DTT; 0.05% Tween80; 5% Glycerol, flash frozen and stored at -80°C.

### ABLE assay

The anti-ORF1p or anti-ORF2p assays are designed to detect anti-ORF1p or anti-ORF2p antibodies using an indirect ELISA method performed in plates coated with ORF1p or a fragment of ORF2p proteins encoded by an L1 retroelement capable of expression in humans. The titers of anti-ORF1p reactive IgG antibodies in human serum were measured in assay plates coated with a full-length ORF1p protein as antigen; the titers of anti-ORF2p reactive IgG antibodies in human serum were measured in assay plates coated with a RT-domain (amino acids residues 367 -771) as antigen as described below.

Nunc MaxiSorp clear 96-well plates were coated with the antigens at 2 µg/mL in phosphate-buffered saline (PBS), 50 µL per well overnight at 4°C. The plates without adsorbed antigen, no-antigen control plates, served to evaluate non-specific antibody binding to plastics.

The assay plates were washed with PBS, containing 0.05% Tween 20 (PBST) then blocked with Blocker Casein in PBS, supplemented with 0.05% Tween 20 (Assay Buffer).

Human serum samples were serially diluted in Assay Buffer, starting with a 100-fold minimum required dilution to a 218700-fold final dilution and were added to the ORF1p, ORF2p and no-antigen control assay plates in duplicates, 50 µL per well. After incubation at room temperature for 2 hours, the assay plates were washed with PBST, and the bound ORF1p- and ORF2p-reactive IgG antibodies were detected with anti-human IgG-HRP (Jackson ImmunoResearch) secondary antibody conjugates diluted 2000-fold in Assay Buffer for 1 hour at room temperature. The plates were washed six times with PBST and incubated with 50 µL of 1-Step Ultra TMB-ELISA Substrate Solution. Peroxidase reactions were stopped with 25 µL 1% hydrochloric acid after 30-minute incubation at room temperature and optical densities at 450 nm (OD450) were measured using a plate reader (Tecan Infinite M1000 Pro). Antibody titers defined as the highest sample dilution that produce positive OD450 reading above a cutpoint value (2-fold OD450 in the wells without test sample) were calculated by a log-linear approximation.

### Anti-p53 antibody ELISA

Recombinant Human Tumor Protein p53 (RayBiotech, Code 230-00639-10) was diluted to 1 µg/mL in a 2M urea/PBS coating buffer and added to Black MaxiSorp 96-well assay plates (50 µL/well) The coating buffer without p53 was added to no-antigen control plates. After overnight incubation at 4°C, assay plates were washed with PBST and blocked with Blocker Casein in PBS, 0.05% Tween 20 (Assay Buffer).

Human serum samples were diluted 300-fold in Assay Buffer and added to the p53 and no-antigen control assay plates, 50 µL/well. After incubation at room temperature for 2 hours, the assay plates were washed with PBST, and the bound p53-reactive IgG antibodies were detected with anti-human IgG-HRP (Jackson ImmunoResearch) diluted 1000-fold in Assay Buffer for 1 hour at room temperature. The plates were washed with PBST and incubated with 50 µL QuantaBlu fluorogenic peroxidase substrate (ThermoFisher) for 15 minutes at room temperature. Fluorescence (RFU) was measured at an excitation wavelength of 320 nm and an emission wavelength of 420 nm using a plate reader (Tecan Infinite M1000 Pro). For each sample, fluorescence measured in no-antigen control wells was subtracted from the corresponding RFU values obtained from p53-coated wells.

### Sandwich ELISA for detection human ORF1p and ORF2p

Capture antibodies: affinity purified rabbit anti-L1 (anti-ORF1p or anti-ORF2p (RT fragment)) polyclonal antibodies were ordered from GenScript (Custom Polyclonal Antibody Production Service). Rabbits were immunized for 7 weeks with human ORF1p protein or human ORF2p peptides. Sera were collected and purified after final immunization using antigen affinity chromatography for ORF1p and protein A affinity chromatography for ORF2p.

Detection antibodies: anti-ORF1p mouse polyclonal sera were obtained in our laboratory by conventional immunization procedure with human ORF1p protein; anti-ORF2p rat affinity purified polyclonal antibodies were obtained from ProSci AbServices after rats ‘ immunization with human ORF2p protein (RT-fragment).

Sandwich ELISA: 50 µL of affinity purified rabbit anti-hORF1p or anti-ORF2p polyclonal antibody immobilized on Black MaxiSorp 96-well assay overnight at 4°C. After washing and blocking with Assay Buffer, the plates were incubated sequentially with serum samples and detection antibodies: anti-ORF1p mouse or anti-ORF2p rat Abs (1:500 in casein assay buffer, 50 µL/well). After washing, goat anti-mouse IgG+IgM+IgA (heavy and light chain) or anti-rat IgG HRP-conjugate in dilution 1:1000 in Assay buffer (50 µL/well) was added and incubated for 1 hour at room temperature, following washing and addition of QuantaBlu fluorogenic peroxidase substrate (ThermoFisher) for 15 minutes at room temperature. Fluorescence was measured at an excitation wavelength of 320 nm and an emission wavelength of 420 nm using a plate reader (Tecan Infinite M1000 Pro).

### Cells with inducible expression of L1

The inducible L1 reporter plasmid used to generate HeLa tet-L1/GLucAI cells (pBH001) was generated through a series of successive PCR-based cloning steps performed by GenScript. pBH001 is comprised of a tetracycline-regulated bidirectional promoter for inducible expression of both firefly luciferase (a fragment cloned from the pTRE3G-BI-Luc control plasmid (Takara Bio)) and a recoded human L1 sequence (encoding both ORF1 and ORF2) cloned from the L1-neo-TET plasmid (Addgene # 51284; a gift from Astrid Roy-Engel)(PubMed 19390602). A custom antisense intron-interrupted *Gaussia* Luciferase (GLuc)-based retrotransposition readout cassette (GLucAI; driven by SV40 promoter) synthesized by GenScript was inserted within the 3 ‘-UTR of L1 in the antisense orientation. Elements originating from the PB-gRNA-puro plasmid (Addgene #121121; a gift from Pablo Navarro) (PubMed 30846691) were used as a backbone for the construct, and includes PiggyBac transposase-specific short inverted terminal repeats (ITRs) (Randolph et al., 2017) that flank the sequence described above and includes a puromycin resistance gene for selection of positive PiggyBac-mediated integration of the cassette into recipient cells.

To generate a stable population of HeLa cells carrying an integrated tetL1-GLucAI construct, HeLa TetOn3G cells (Takara) were co-transfected with the tet-L1/GLucAI donor plasmid (pBH201) and the Super PiggyBac Transposase Expression Vector (System Biosciences) at a 5:1 plasmid ratio, as recommended by the manufacturer’s protocol. Transfections were performed using LipoD293 reagent (SignaGen) with a total of 3 μg plasmid DNA per 10-cm dish, following the manufacturer’s protocol. 72h after transfection, cells were selected for 3 days in 1 μg/mL puromycin. After 2 passages, an additional 72h selection was performed to ensure acquisition of a population with stably integration of the tetL1-GLucAI cassette. Clones were isolated via limiting dilution into 96-well plates, and subsequently characterized for doxycycline-stimulated acquisition of *Gaussia* luciferase activity. Clone 9A8 was used in this study.

### Retrotransposition assay

HeLa tet-L1/GLucAI were plated at 10^6^ cells per 10-cm plate and expression of the integrated tet-L1/GLucAI cassette was induced by addition of doxycycline (Dox) to a final concentration of 400 ng/mL in the culture medium. After 48 hours, GLuc activity in the conditioned medium (indicative of successful L1-mediated retrotransposition activity) was measured using a microplate luminometer immediately following addition of an equal volume of 2x GLuc reagent (50 μM coelenterazine (GoldBio) in D-PBS containing 300 mM sodium ascorbate (Sigma) and 0.2% Triton X-100 (Sigma)). HeLa tet-L1/GLucAI cells without induction were used as control and prepared by the same way (**Fig. S2**).

### Western immunoblotting

Plates with HeLa tet-L1/GLucAI cells +/-Dox were placed on ice and the cells were washed with ice-cold PBS. After PBS aspiration, cells were sonicated in ice cold RIPA buffer (Sigma) plus protease inhibitors (PIC) (Sigma) and cleared by centrifugation in a Beckman tabletop (20-minute, 12,000 rpm, 4°C). The protein concentrations of lysates were determined with the Quick start Bradford reagent (Bio-Rad), using bovine serum albumin (BSA) (Sigma) as a standard. Cells lysates were separated by BioRad mini protean TGX gels 4-20%, 15 μg of total cell protein were loaded per well. Precision plus dual color protein molecular weight markers (Bio-Rad) were also loaded. For Western blots, proteins were then transferred to Immun-Blot PVDF membranes (Bio-Rad). After blocking in 2.5% non-fat dry milk (RPI) in TBST (10 mM Tris-HCl (ThermoFisher) pH 7.5, 150 mM NaCl (ThermoFisher), 0.1% Tween-20 (Sigma)) for 1 hour at room temperature, blots were incubated with primary antibodies against ORF1p overnight at 4°C with constant shaking: 1) anti-ORF1p IgG high-titers human serum samples in 1% non-fat dry milk (RPI) TBST in dilution 1:1,000-1000,000, followed by horseradish peroxidase (HRP)-conjugated secondary anti-human IgG antibodies (1:10,000 dilution, Sera Care) 1 hour at room temperature or 2) rabbit anti-human ORF1p polyclonal serum in 1% non-fat dry milk (RPI) in TBST in dilution 1:1,000, followed by horseradish peroxidase (HRP)-conjugated secondary anti-mouse IgG antibodies (1:4,000 dilution, Santa Cruz) 1 hour at room temperature. Membranes were washed three times with TBST, then developed by ECL (Perkin-Elmer) and documented using the ChemiDoc MP Imaging System (BioRad).

### Immunofluorescence staining of cells

Cells were fixed in 4% formaldehyde in PBS for 15 minutes at 4°C, washed 3x in PBS, kept at 4°C for 1 hour in PBS for staining, blocked in blocking solution (5% normal donkey serum, 0.2% triton x-100, PBS or TBS, 1% glycine) for 15 minutes. Cells were incubated with primary antibody (mouse monoclonal from Millipore, MABS1152 dilution 1:400 in block solution or human anti-ORF1p antibody (#11, #766, #899, #865 1:50,000 dilution or rabbit anti-human ORF1p custom made 1:200 dilution for 30-60 minutes, washed 6x in PBS, cells were incubated with secondary antibody (donkey anti-mouse, anti-human or anti-rabbit Jackson ImmunoResearch laboratories Inc) Cy3 or AlexaFluor488 conjugated, 2µg/mL for 30 minutes, washed 6x in PBS. Nuclei were counterstained with DAPI. Samples were washed 4 hours with several changes of PBS, cleared, and mounted with ProLong Gold antifade reagent. Images were collected with Zeiss AxioImager 2 fluorescent microscope equipped with AxioCam 702 digital camera using ZEN software 2.6 version.

### Statistical analysis

Statistical analysis and graph plotting were performed using GraphPad Prizm software (Version 9) and Microsoft Office Excel. To determine the normality of the distribution the Shapiro-Wilk normality test was used. The differences in multiple groups were analyzed by Dunn’s multiple comparison test with the multiplicity adjusted p-value. The differences between two groups were analyzed by the Mann– Whitney U-test. For correlation analysis Spearman’s R-correlation analysis was performed. Logistic regression was performed with disease status as response and anti-ORF1p IgG as independent variable, while adjusting for age. A value of p<0.05 was considered statistically significant in all analyses.

