## Supplemental Information for "Cancer relevance of circulating antibodies against LINE-1 antigens in humans"

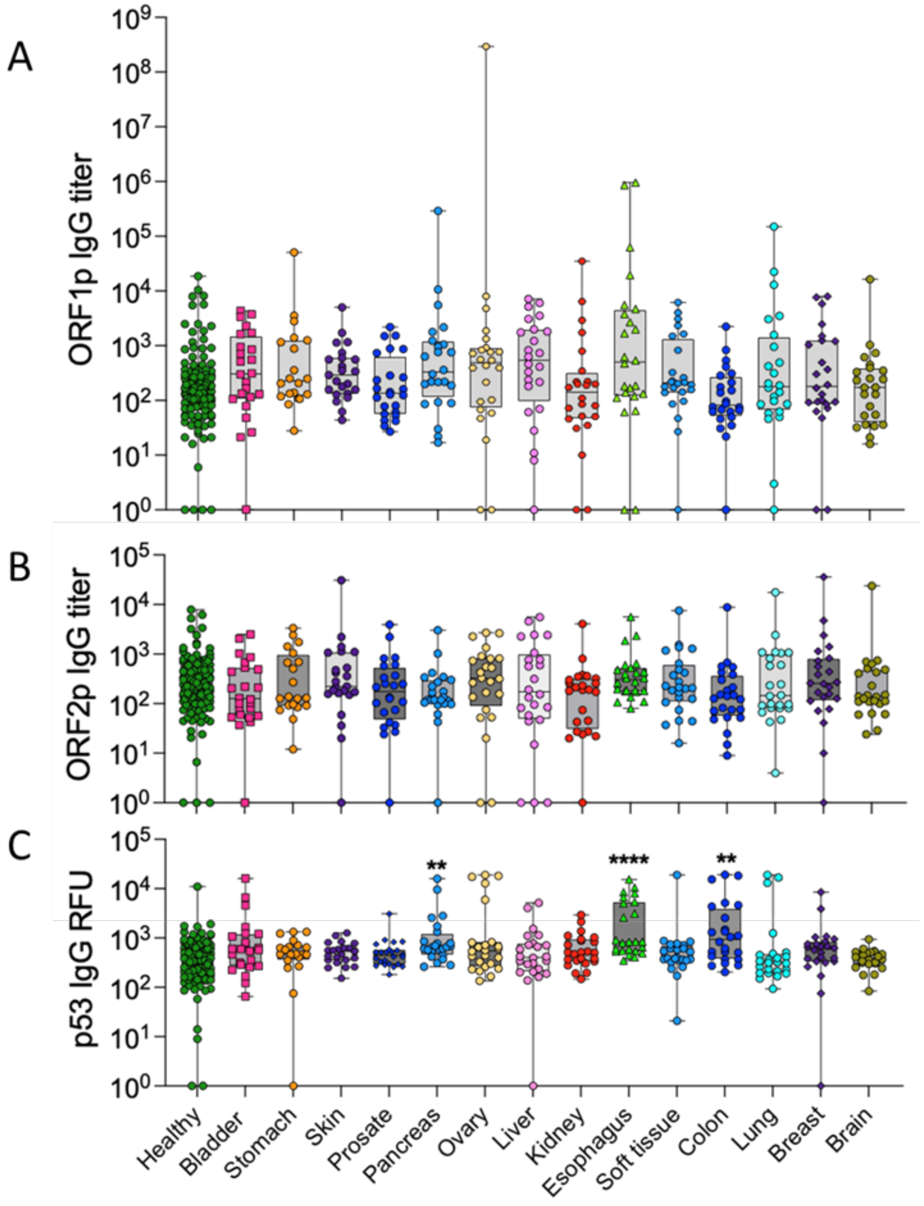


**Figure S1. Anti-ORF1p, anti-ORF2p IgG titers and anti-p53 IgG signals in serum samples of patients with 14 cancer types and healthy individuals. A.** Boxplots for anti-ORF1p IgG titers, **B.** anti-ORF2p IgG titers and **C.** anti-p53 IgG signals represented by median with range and individual values. Statistics were calculated by Dunn's multiple comparison test with adjusted p-value for anti-ORF1p, anti-ORF2p IgG titers and anti-p53 IgG signals in ELISA for serum samples representing 14 solid cancer types: ovarian (N=24), breast (N=24), lung cancer (N=24), colorectal (N=24), esophageal (N=24), renal (N=24), liver (N=24), pancreatic (N=24), prostatic (N=24), gastric (N=20), bladder (N=24) cancer, soft tissue sarcoma (N=24), melanoma (N=23), glioblastoma (N=24), vs. healthy individuals (N=137). **, P < 0.01; ****, P < 0.0001.

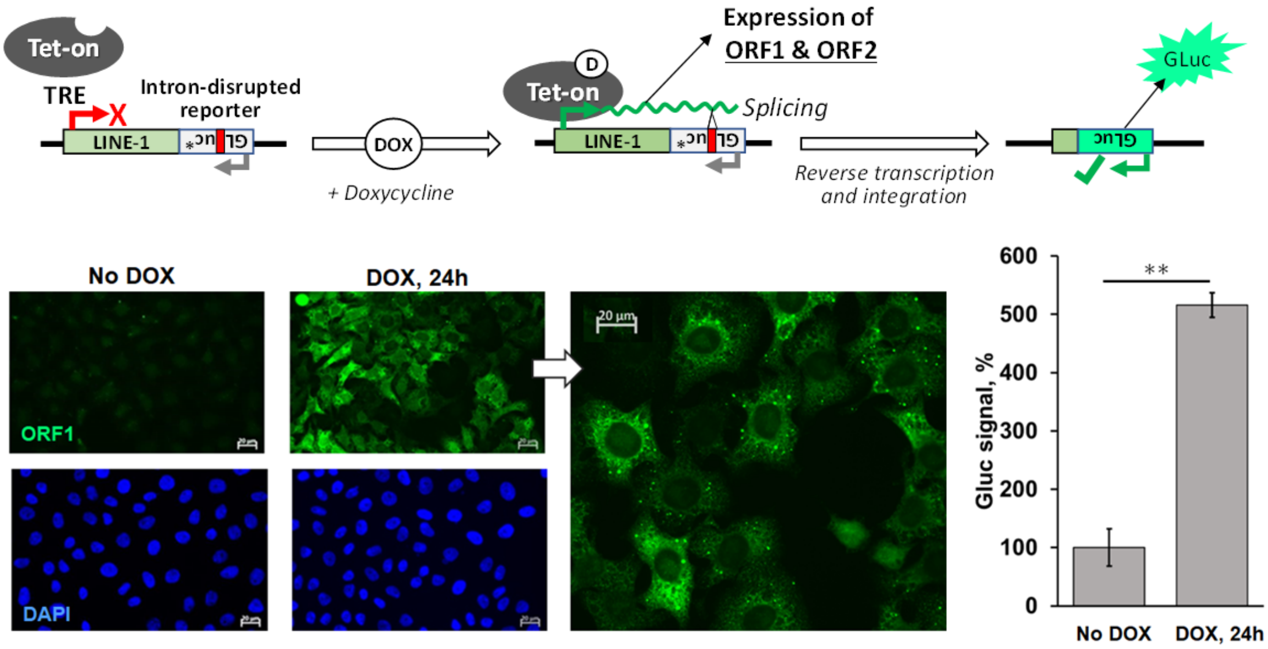


**Figure S2. Characterization of HeLa tet-L1/GLucAI cells.** A schematic of the doxycycline-inducible L1 cassette stably integrated into HeLa cells (top panel). Fluorescent microphotographs depict the induction of human ORF1 in HeLa tet-L1/GLucAI cells following a 48-hour induction with 100 ng/mL doxycycline, as demonstrated by punctate cytoplasmic immunoreactivity by anti-human ORF1 antibodies (green) with DAPI nuclear counterstain (blue) (bottom left). Following a 48-hour exposure to doxycycline (400 ng/mL), an induction of secreted *Gaussia* luciferase activity (a readout for L1-mediated retrotransposition) was observed (bottom right). Mean and standard deviation of normalized data (compared to non-induced control) are depicted. Statistics were calculated by Student's t-test (unpaired, two-tailed); ** p-value < 0.01.

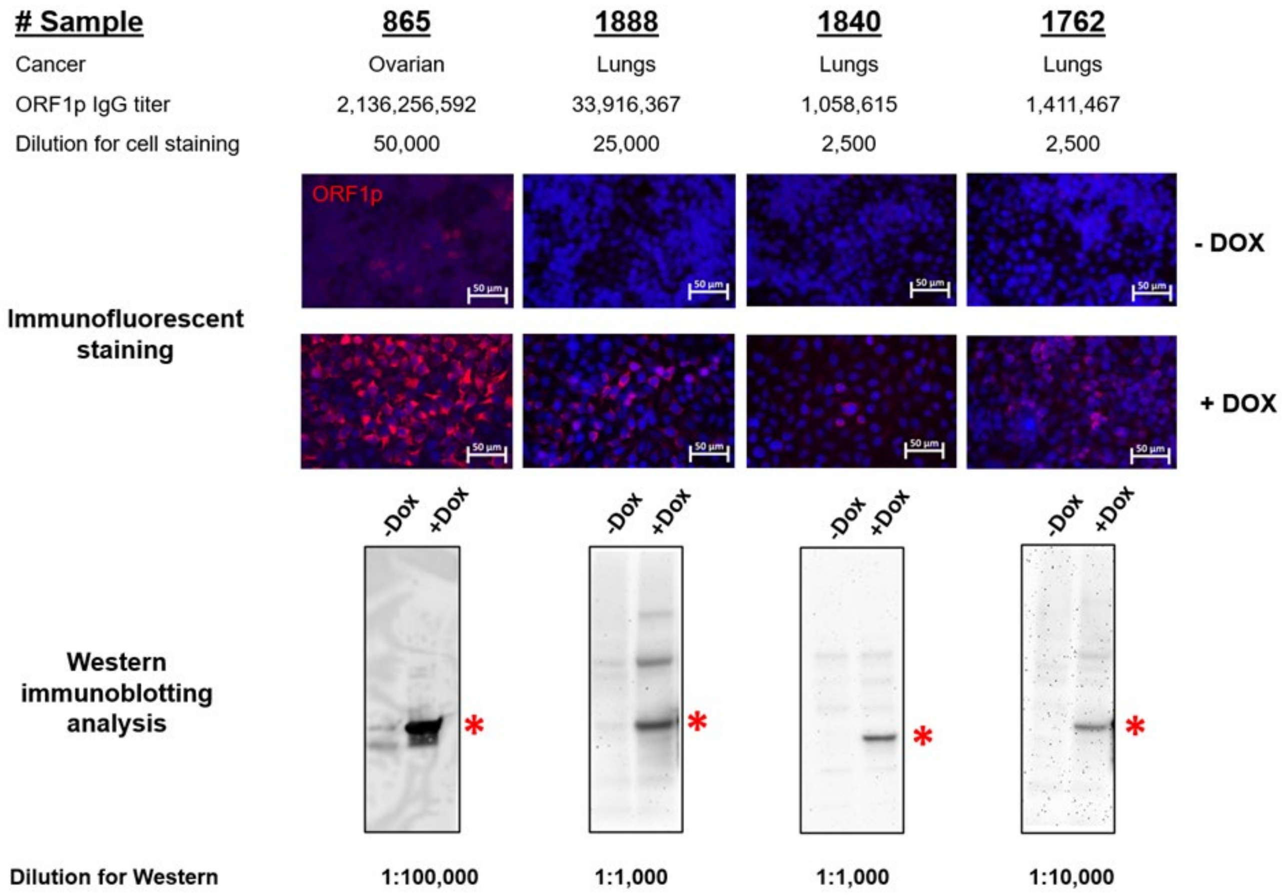


**Figure S3. Relevance of antibodies detected by ABLE to ORF1p.** The results of immunofluorescent staining (upper panel) and immunoblot analysis (lower panel) of HeLa cells transduced with Tet-inducible L1/GlucAI reporter construct with (+ Dox) and without (- Dox) L1 induction by doxycycline for 24 hours. The panels represent staining with diluted serum samples from ovarian and lung cancer patients scored positive in ABLE$^{ORF1}$ assay (#865, #1888, #1840, #1762). The asterisk on the lower panel corresponds to 40-kDa band of ORF1p recombinant protein standard.

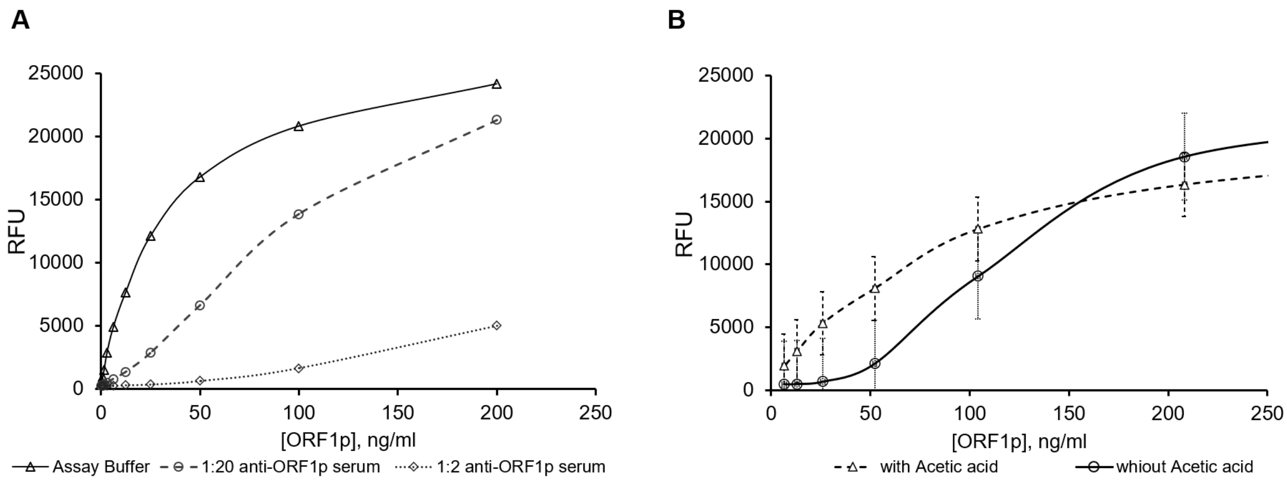


**Figure S4. Detection of human ORF1p antigen by sandwich ELISA in the presence of human anti-ORF1p-reactive antibodies. A.** Concentration dependence of anti-ORF1p-reactive antibodies from human serum on the detection of human ORF1p antigen standard curve by sandwich ELISA. **B.** Human ORF1p antigen calibration plot in serum with added human anti-ORF1p reactive antibodies with and without acid dissociation detected by sandwich ELISA.

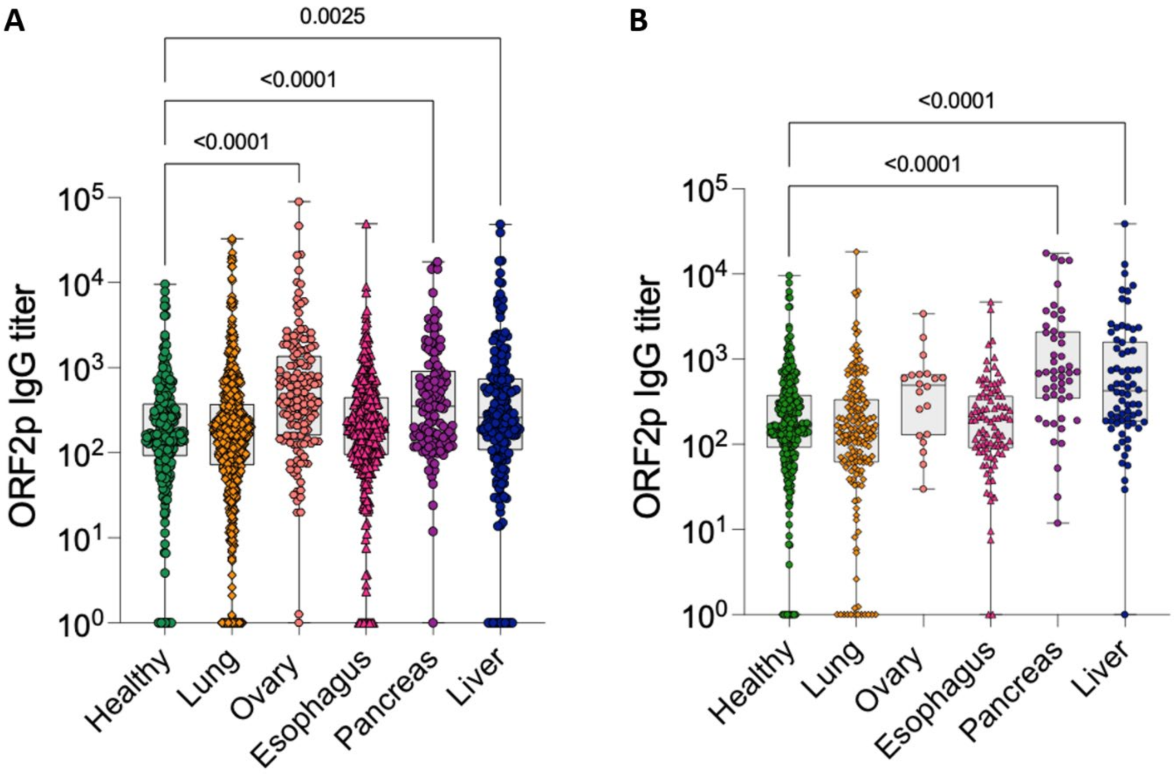


**Figure S5. Analysis of anti-ORF2p IgG titers in esophageal, lung, pancreatic, ovarian and liver cancer. A.** Detection of anti-ORF2p IgG titers in serum samples of patients with indicated cancer types (all stages). Sample size: healthy (N=274), lung (N=707), ovary (N=150), esophagus (N=377), pancreas (N=124), and liver (N=217) **B.** The same as panel A, except only samples from patients with cancer stages 1 and 2 are shown. Sample size: healthy (N=274), lung (N=169), ovary (N=20), esophagus (N=92), pancreas (N=48), and liver (N=70). Boxplots for ORF2p IgG titers for five selected cancer types depicting median with range and individual values. Statistics were calculated by Dunn's multiple comparison test with multiplicity-adjusted p-values for anti-ORF2p IgG titers in ELISA for five cancer categories vs. healthy individuals. All p-values < 0.05 are indicated.

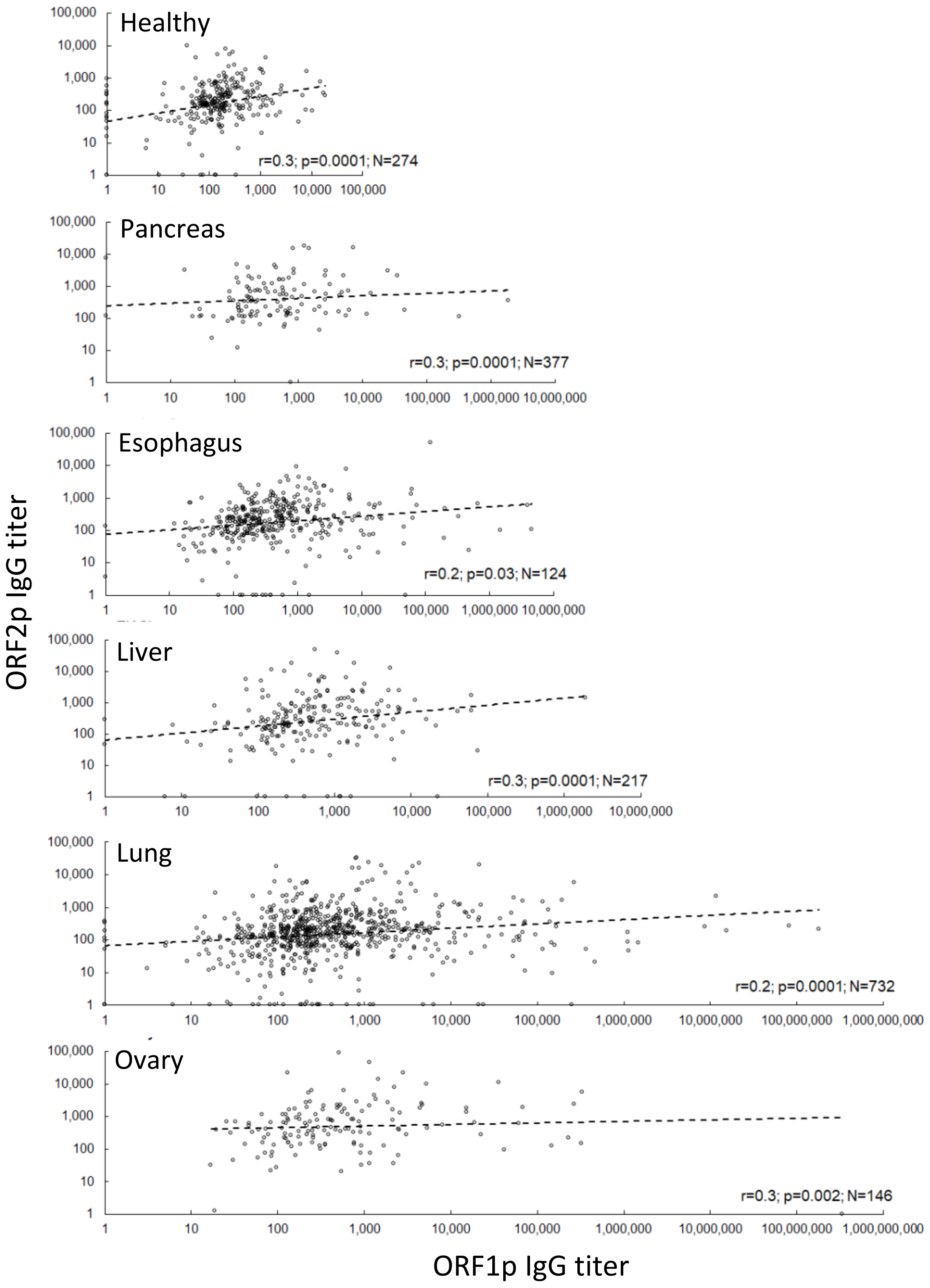


**Figure S6.** Spearman's correlation analysis of anti-ORF1p and anti-ORF2p IgG titers in patients with five cancer types and in healthy individuals.

A

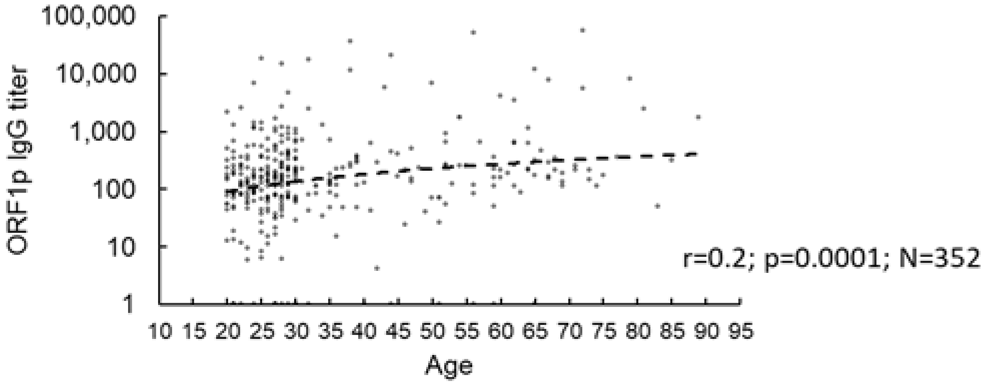


B

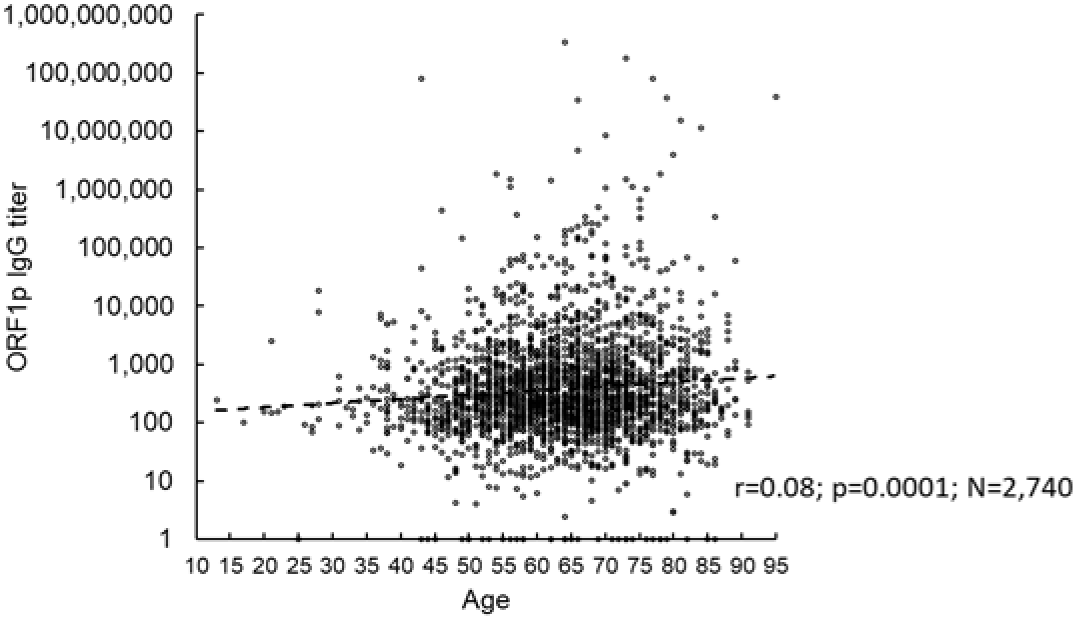


**Figure S7.** Spearman's R-correlation analysis of anti-ORF1p IgG titers and age among A. healthy (20-89 years of age) or B. Cancer patients (13-95 years of age).

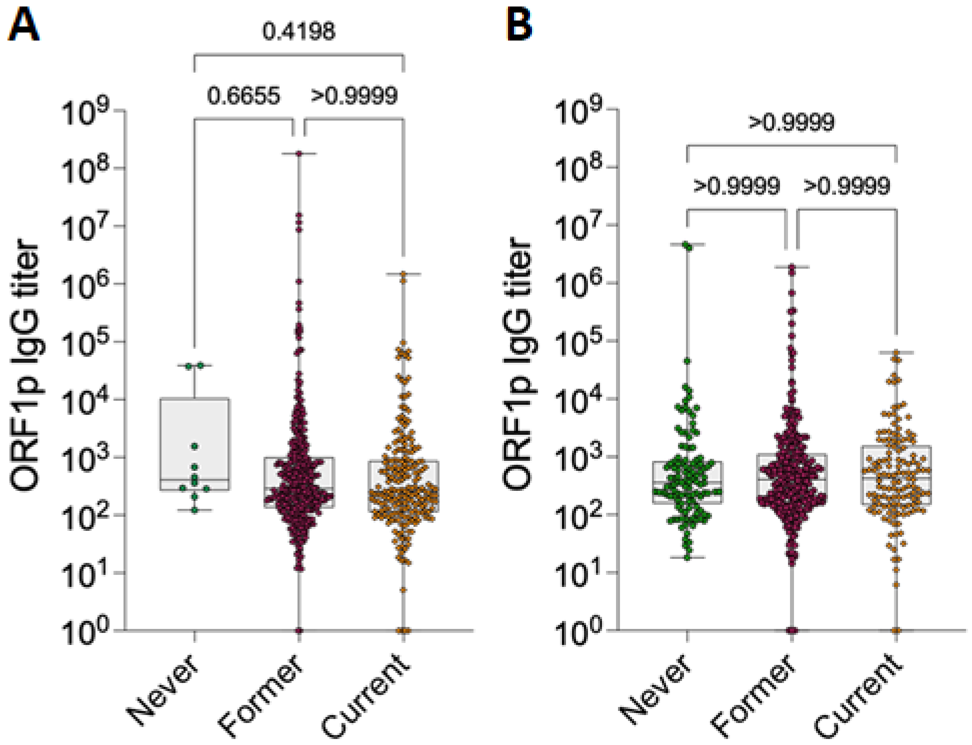


**Figure S8. Anti-ORF1p IgG titers in tobacco smokers' groups for lung cancer patients (A) and for cancer patients from three cancer categories (pancreatic, esophageal and liver) (B).** A. Lung cancer: former smokers (N=321), never smokers (N=10), current smokers (N=205). B. Combined cancer group: former smokers (N=266), never smokers (N=115), current smokers (N=133). Boxplots for anti-ORF1p IgG titers for patients with lungs cancer grouped by smoking status, depicting the minimum, first quartile, median, third quartile, maximum and individual titer values. Statistics were calculated by Dunn's multiple comparison test with adjusted p-values for anti-ORF1p IgG titers determined by ELISA for three groups: former smokers, current smokers, and never smokers. All p-values > 0.05.

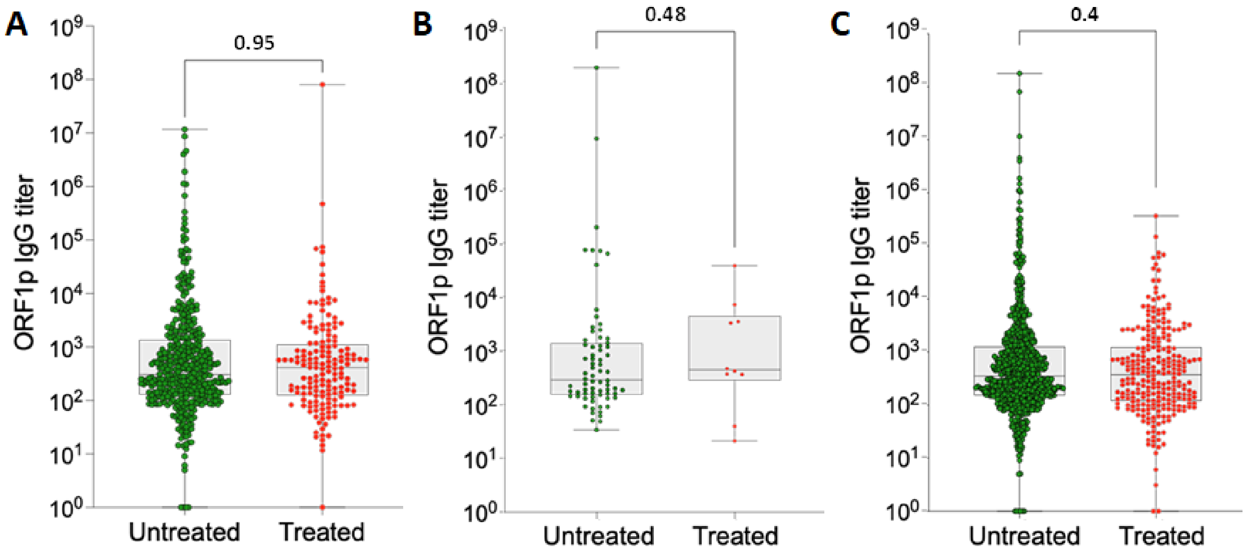


**Figure S9. Cross-sectional study of cancer patients with or without exposure to anti-cancer therapies.** Anti-ORF1p IgG titers in cancer patients combined from four cancer types (lungs, pancreatic, esophageal, and liver). **A.** Radiation therapy: untreated (N=442), treated (N=161). **B.** Immunotherapy: untreated (N=72), treated (N=10). **C.** Chemotherapy: untreated (N=771), treated (N=266). Boxplots for anti-ORF1p IgG titers combined for patients with four cancer types grouped by therapy, depicting the minimum, first quartile, median, third quartile, maximum and individual titer values. Statistics were calculated by Mann-Whitney test for anti-ORF1p IgG titers determined by ELISA for treated vs. untreated cancer patients. All p-values > 0.05.

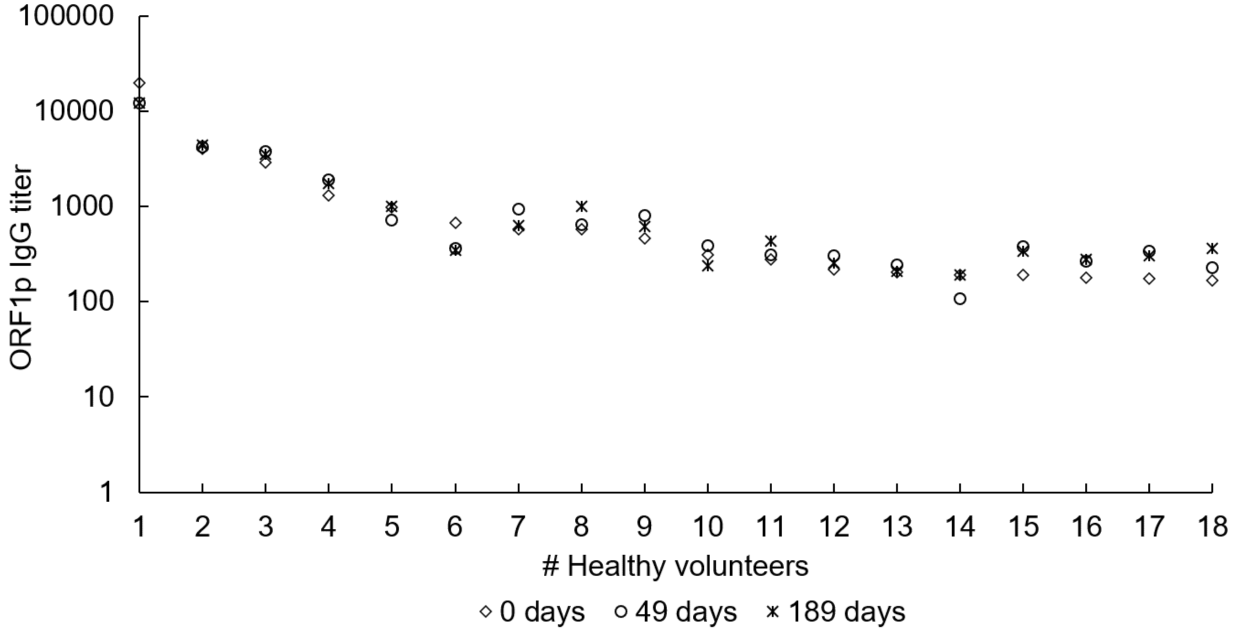


**Figure S10. Longitudinal study of anti-ORF1p IgG titers in serum samples from healthy volunteers.** Blood was drawn from 18 healthy individuals (66±10 (mean±SD) years of age) on day 0, 49, and 189. Anti-ORF1 IgG titers were determined by ELISA. The average coefficient of variation for titer measurements from individual volunteers was 25%.

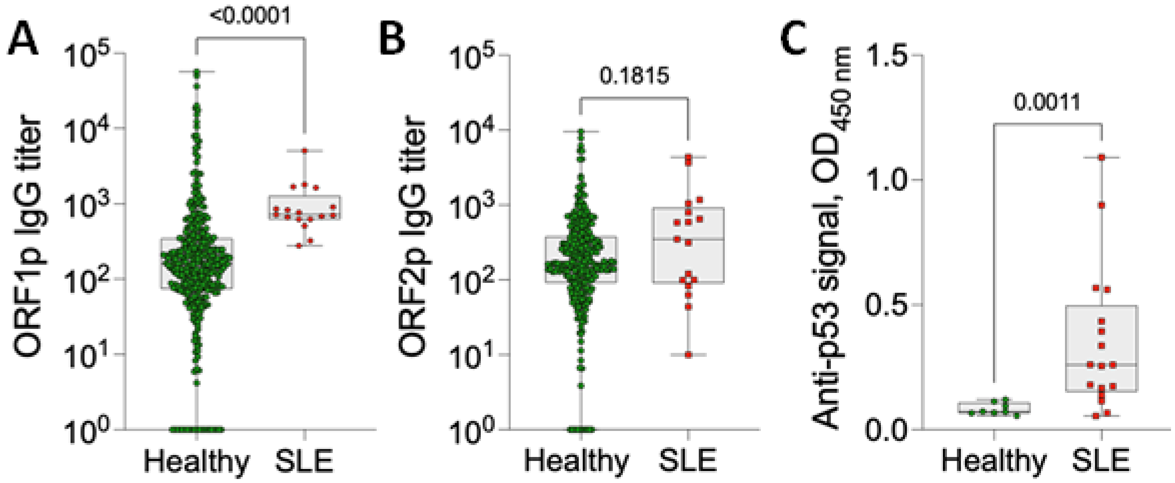


**Figure S11. Analysis of anti-ORF1p, anti-ORF2p and anti-p53 IgG in blood samples from patients with SLE and healthy individuals. A.** Anti-ORF1p IgG titers in samples of SLE patients (N=17) vs. healthy control samples (N=352) determined by ELISA. **B.** Anti-ORF2p IgG titers in samples of SLE patients (n=17) vs. healthy control (N=274) determined by ELISA. **C.** Anti-p53 signals in **"**p53 Autoantibody ELISA Kit (Human)" (Dianova). Statistics were calculated by Mann-Whitney U-test, p-value < 0.05 is considered significant.

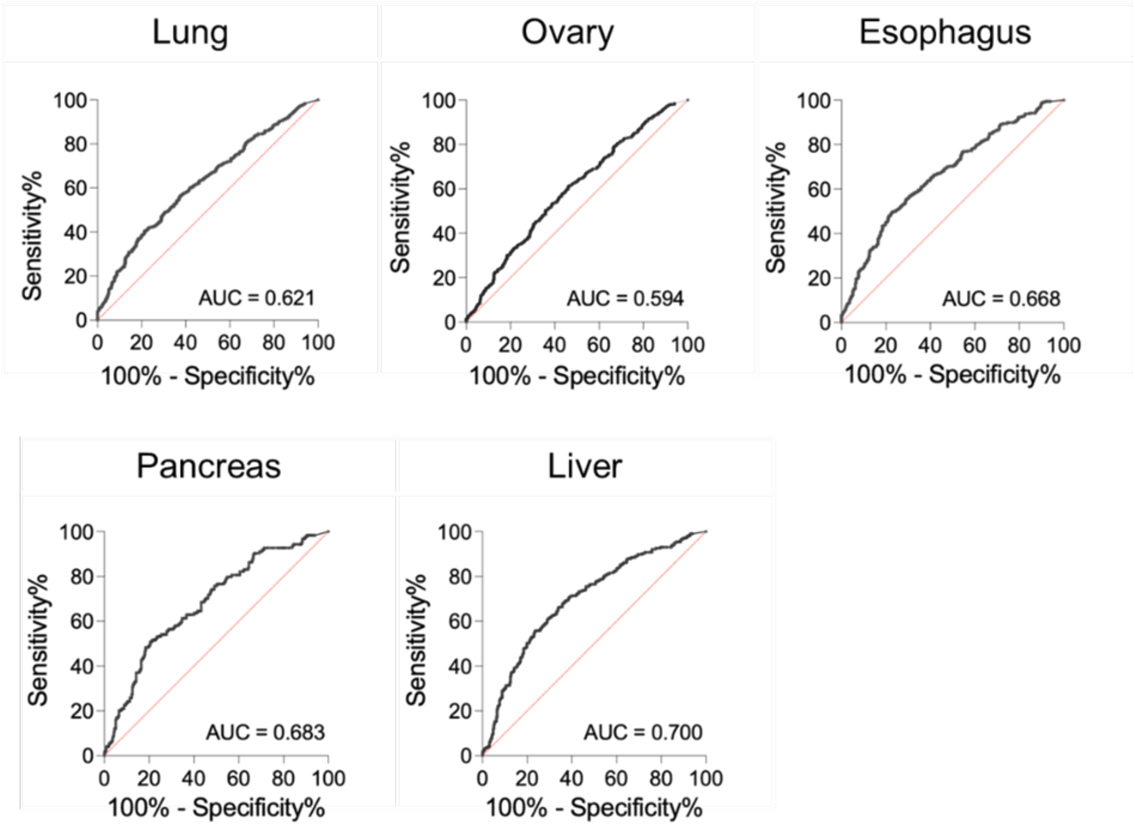


**Figure S12. The ROC curves indicating specificity and sensitivity of ABLE$^{ORF1}$ for 5 cancer types.** Lung (N=908), ovary (N=993), esophagus (N=377), pancreas (N=124), liver (N=217) cancers and healthy (N=352). All p-values < 0.0001.

### Supplementary Tables

**Table S1. Anti-ORF1p IgG titers detected in ABLE and Western blot (WB)-based detection of ORF1p.** Detection of 40-kDa band on WB (from doxycycline-induced HeLa tet-L1/GLucAI cells or human ORF1p recombinant standard) stained by anti-ORF1p human IgG from the serum samples of cancer patients and healthy volunteers.

| # Sample | Cancer type | ORF1p IgG titers | ORF1p on WB |
|---|---|---|---|
| 24 | **Healthy** | 18,634 | - |
| 184 | | 17,584 | - |
| 76 | | 15,209 | - |
| 134 | | 10,491 | + |
| 322 | | 8,166 | - |
| 131 | | 7,943 | - |
| 69 | | 1,321 | - |
| 185 | | 181 | - |
| 218 | | 137 | - |
| 96 | | 49 | - |
| 1510 | **Lung** | 180,046,944 | + |
| 1206 | | 80,557,768 | + |
| 1888 | | 33,916,367 | + |
| 1390 | | 11,669,656 | + |
| 373 | | 5,320,280 | + |
| 1132 | | 1,482,912 | + |
| 1762 | | 1,411,483 | + |
| 1376 | | 1,129,245 | + |
| 1050 | | 1,103,532 | - |
| 1840 | | 1,058,615 | + |
| 1189 | | 1,045,443 | + |
| 1025 | | 469,697 | + |
| 990 | | 369,076 | + |
| 384 | | 269,281 | - |
| 1174 | | 253,275 | + |
| 1294 | | 195,648 | + |
| 1071 | | 169,520 | + |
| 1071 | | 169,520 | + |
| 1080 | | 149,600 | + |
| 297 | | 149,257 | - |
| 1254 | | 134,583 | + |
| 1884 | | 71,173 | - |
| 1409 | | 20,283 | - |
| 1892 | | 19,236 | + |
| 535 | | 17,367 | + |
| 1364 | | 15,271 | - |
| 541 | | 15,132 | + |

| | | | |
|---|---|---|---|
| 156 | | 12,947 | + |
| 1663 | | 11,289 | + |
| 1615 | | 10,245 | - |
| 865 | **Ovary** | 462,671,117 | + |
| 425 | | 293,944,784 | + |
| 766 | | 81,119,792 | + |
| 899 | | 39,170,713 | + |
| 11 | | 37,500,953 | + |
| 859 | | 438,572 | + |
| 559 | | 329,307 | + |
| 1-48 | | 327,029 | + |
| 2-18 | | 292,580 | + |
| 2-62 | | 283,251 | + |
| 2-29 | | 275,814 | + |
| 1-05 | | 164,614 | + |
| 916 | | 103,501 | + |
| 1-31 | | 12,005 | - |
| 240 | | 883 | + |
| 4 | | 859 | - |
| 195 | | 251 | - |
| 896 | | 212 | - |
| 193 | **Esophagus** | 960,314 | + |
| 149 | | 856,547 | + |
| 973 | | 672,893 | + |
| 1092 | | 493,030 | - |
| 374 | | 121,991 | + |
| 312 | | 63,661 | + |
| 886 | **Liver** | 1,891,171 | + |
| 772 | | 40,371 | - |
| 719 | **Pancreas** | 1,871,054 | - |
| 421 | | 290,552 | + |

**Table S2.** Logistic regression results for anti-ORF1 IgG titers in cancer and healthy after adjusting for age.

| | Estimate | Std. Error | z value | p-value |
|---|---|---|---|---|
| **Anti-ORF1p IgG titer** | 0.2305 | 0.1051 | 2.1941 | 0.0282 |
| **Age** | 0.1401 | 0.0061 | 22.8113 | <0.0001 |

**Table S3. Comparison of anti-ORF1p IgG titers among healthy individuals.** Dunn's multiple comparison test with adjusted p-value; p-value < 0.05 is considered significant.

| Ethnicity (sample size) | Comparison | p-value |
|---|---|---|
| Black (N=37) | Black vs. Caucasian | p=0.9 |
| Hispanic (N=143) | Hispanic vs. Black | p=0.8 |
| Caucasian (N=167) | Caucasian vs. Hispanic | p=0.3 |

**Table S4. Gender-specificity of anti-ORF1p IgG titers among healthy individuals and cancer patients.** Mann-Whitney U-test, p-value < 0.05 is considered significant.

<table>
<tr><th>Cancer type</th><th>Sample size</th><th>p-value</th></tr>
<tr><td rowspan="2">Healthy</td><td>Male (N=159)</td><td rowspan="2">p=0.2</td></tr>
<tr><td>Female (N=188)</td></tr>
<tr><td rowspan="2">Pancreas</td><td>Male (N=52)</td><td rowspan="2">p=0.3</td></tr>
<tr><td>Female (N=72)</td></tr>
<tr><td rowspan="2">Liver</td><td>Male (N=185)</td><td rowspan="2">p=0.4</td></tr>
<tr><td>Female (N=32)</td></tr>
<tr><td rowspan="2">Esophagus</td><td>Male (N=320)</td><td rowspan="2">p=0.5</td></tr>
<tr><td>Female (N=57)</td></tr>
<tr><td rowspan="3">Lung</td><td>Male (N=451)</td><td rowspan="3">p=0.004</td></tr>
<tr><td>Female (N=456)</td></tr>
<tr><td>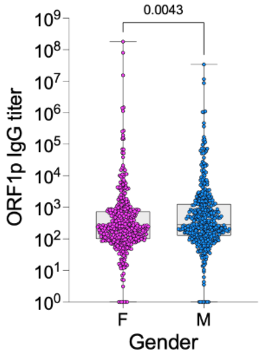
</td></tr>
</table>

**Table S5. Anti-ORF1p IgG titers in subjects with myocardial infarction (MI) vs. healthy or chronic obstructive pulmonary disease/pneumonia (COPD/PN) patients vs. healthy and lung cancer patients.** Mann-Whitney U-test, p-value < 0.05 is considered significant.

| **Anti-ORF1p IgG titers** | **p-value** |
|---|---|
| COPD/PN patients (N=37) ~ healthy (n=304) | 0.2 |
| COPD/PN patients (N=37) **<** lung cancer patients (N=708) | 0.0004 |
| MI patients (N=33) ~ healthy (N=304) | 0.9 |

**Table S6.**

| **Cancer type** | **Sample size** | | |
|---|---|---|---|
| | **All samples** | **Stages 1-2** | **Stages 3-4 and others** |
| **Ovary** | N=979 | N=193 | N=786 |
| **Pancreas** | N=124 | N=40 | N=84 |
| **Liver** | N=217 | N=67 | N=150 |
| **Esophagus** | N=377 | N=79 | N=298 |
| **Lung** | N=907 | N=90 | N=817 |
